# Chromosomal fusion and life history-associated genomic variation contribute to within-river local adaptation of Atlantic salmon

**DOI:** 10.1101/347724

**Authors:** Kyle Wellband, Claire Mérot, Tommi Linnansaari, J. A. K. Elliott, R. Allen Curry, Louis Bernatchez

## Abstract

Chromosomal inversions have been implicated in facilitating adaptation in the face of high levels of gene flow, but whether chromosomal fusions also have similar potential remains poorly understood. Atlantic salmon are usually characterized by population structure at multiple spatial scales; however, this is not the case for tributaries of the Miramichi River in North America. To resolve genetic relationships between populations in this system and the potential for known chromosomal fusions to contribute to adaptation we genotyped 728 juvenile salmon using a 50K SNP array. Consistent with previous work, we report extremely weak overall population structuring (Global F_ST_ = 0.004) and failed to support hierarchical structure between the river’s two main branches. We provide the first genomic characterization of a previously described polymorphic fusion between chromosomes 8 and 29. Fusion genomic characteristics included high LD, reduced heterozygosity in the fused homokaryotes, and strong divergence between the fused and the unfused rearrangement. Population structure based on fusion karyotype was five times stronger than neutral variation (F_ST_ = 0.019) and the frequency of the fusion was associated with summer precipitation supporting a hypothesis that this rearrangement may contribute local adaptation despite weak neutral differentiation. Additionally, both outlier variation among populations and a polygenic framework for characterizing adaptive variation in relation to climate identified a 250 Kb region of chromosome 9, including the gene *six6* that has previously been linked to age-at-maturity and run-timing for this species. Overall our results indicate that adaptive processes, independent of major river branching, are more important than neutral processes for structuring these populations.

## Introduction

Conservation and management of species benefit from an understanding of fine-scale demographic and evolutionary processes, which aid in delimitating appropriate population-scale units for management and conservation of genetic diversity (Fraser & Bernatchez, 2001; Funk, McKay, Hohenlohe, & Allendorf, 2012; Palsbøll, Bérubé, & Allendorf, 2007; Waples & Gaggiotti, 2006). Population genomic methods provide increased power for delineating genetic structure over traditional approaches and can identify putatively adaptive variation among populations that may reflect local adaptation (Funk et al., 2012). Genomic variation associated with environmental factors can indicate relevant biotic or abiotic forces possibly driving local adaptation (e.g. Benestan et al., 2016; Bourret, Dionne, Kent, Lien, & Bernatchez, 2013; Hecht, Matala, Hess, & Narum, 2015). Furthermore, the genomic basis of important life history variation relevant to maintaining within-population diversity can be identified (Barson et al., 2015; Cauwelier, Gilbey, Sampayo, Stradmeyer, & Middlemas, 2017). Knowledge of genomic variation linked to various locally-adaptive phenotypes will facilitate improved population monitoring and more effective conservation and management of species (Aykanat, Lindqvist, Pritchard, & Primmer, 2016).

Population genomics has also helped reveal the role of chromosomal rearrangements in eco-evolutionary processes (Wellenreuther & Bernatchez, 2018). Large rearrangements (e.g. chromosomal inversions, fusions, fissions, or translocations) disturb homologous chromosomal pairing during meiosis that results in tight linkage among genes within rearranged regions detectable by genomic methods (Wellenreuther & Bernatchez, 2018). Chromosomal rearrangements function like “supergenes” and have been implicated in important adaptive phenotypes of several species (e.g., salinity in cod: Berg et al., 2015, migratory behaviour in cod: Berg et al., 2016; climatic gradients in Drosophila: Kapun, Fabian, Goudet, & Flatt, 2016; crypsis in stick insects: Lindtke et al., 2017). Furthermore, they have been shown to facilitate environmental adaptation in the face of high levels of gene flow (Barth et al., 2017). Chromosomal fusions generate karyotype variation between as well as within species (Dobigny, Britton-Davidian, & Robinson, 2017). While a potential role for chromosomal fusion in facilitating local adaptation has been demonstrated theoretically (Guerrero & Kirkpatrick, 2014), empirical evidence for this phenomenon is lacking.

Atlantic salmon, *Salmo salar*, is an economically and culturally important anadromous fish on both sides of the Atlantic Ocean. The species exhibits a wide variety of life history strategies, including the timing of migrations and differing ages at maturity, that are present throughout large portions of its range and are believed to reflect local adaptation to a variety of biotic and abiotic conditions (e.g. temperatures, stream flow, pathogens, and predators; Fraser, Weir, Bernatchez, Hansen, & Taylor, 2011; Garcia de Leaniz et al., 2007; Riddell & Leggett, 1981; Riddell, Leggett, & Saunders, 1981; Schaffer & Elson, 1975). A combination of natal philopatry and low rates of straying result in structuring where river populations are differentiated from one another but cluster into regional groups. This hierarchical population structure occurs in both Europe and North America (Bourret, Kent, et al., 2013; Dionne, Caron, Dodson, & Bernatchez, 2008; Perrier, Guyomard, Bagliniere, & Evanno, 2011; Verspoor et al., 2005). Regional structure is driven by a combination of geographic proximity of individual populations as well as climatic gradients (e.g. temperature, precipitation) suggesting that adaptive processes are also important for structuring regional gene pools (Bourret, Dionne, et al., 2013; Dionne, Miller, Dodson, Caron, & Bernatchez, 2007; Jeffery et al., 2017). At finer scales, population structure can be present among tributaries within certain river systems (Aykanat et al., 2015; Cauwelier et al., 2017; Dionne, Caron, Dodson, & Bernatchez, 2009; Garant, Dodson, & Bernatchez, 2000; Primmer et al., 2006; Pritchard et al., 2018; Vähä, Erkinaro, Niemelä, & Primmer, 2007). Within-river structuring is also believed to reflect local adaptation because even weak genetic structure within river systems can correspond to biologically meaningful differences in growth and survival among populations (e.g. Aykanat et al., 2015; Mobley et al., 2018). Despite the belief that Atlantic salmon exhibit widespread local adaptation, the genomic basis of many adaptive traits remains unclear. Furthermore, Atlantic salmon from North America are known to be polymorphic for multiple chromosomal fusions (Brenna-Hansen et al., 2012), but their variability in natural populations and potential role in adaptation remain unknown.

The Miramichi River, New Brunswick, Canada, has a large watershed with a complex river network with no unnatural barriers for fish movement (Cunjak & Newbury, 2005). Despite harbouring a large population of Atlantic salmon with some evidence of demographic independence between at least the two main branches of the river (DFO, 2017), several studies have failed to reveal significant genetic population structure in this system (Dionne et al., 2009; Dodson & Colombani, 1997; Moore et al., 2014; but see Møller, 2005). These studies were based on limited numbers of neutral markers and modest sampling designs, leaving uncertainties about the source of negative results (panmixia) and without the possibility of inferring adaptive variation. Here we revisit this question and employ both an extensive sampling design and high-resolution genotyping to characterize spatial patterns of within-river population structure, the potential for chromosomal fusions to facilitate adaptation despite low levels of genetic structure, and identify spatial patterns of putatively adaptive differentiation for Atlantic salmon in the Miramichi River. This work contributes to our knowledge of the patterns and processes of local adaptation in large river systems.

## Methods

### Sampling and SNP array genotyping

Juvenile Atlantic salmon were collected by electroshocking from sites throughout the Miramichi River watershed from August 29 to October 13, 2016 (Figure 1, Table S1). Samples were humanely euthanized and transported on ice to the Miramichi Salmon Association Conservation Rearing Facility at South Esk, New Brunswick. Age 1+juveniles were identified based on established length-age relationships (Swansburg, Chaput, Moore, Caissie, & El-Jabi, 2002) and fin clips were removed and preserved in 95% ethanol. We assessed the modality of lengths and weights in each tributary using Hartigan’s dip test for modality (Hartigan & Hartigan, 1985) and corrected for multiple tests using a false discovery rate (FDR) correction (Benjamini & Hochberg, 1995).

**Figure 1:**
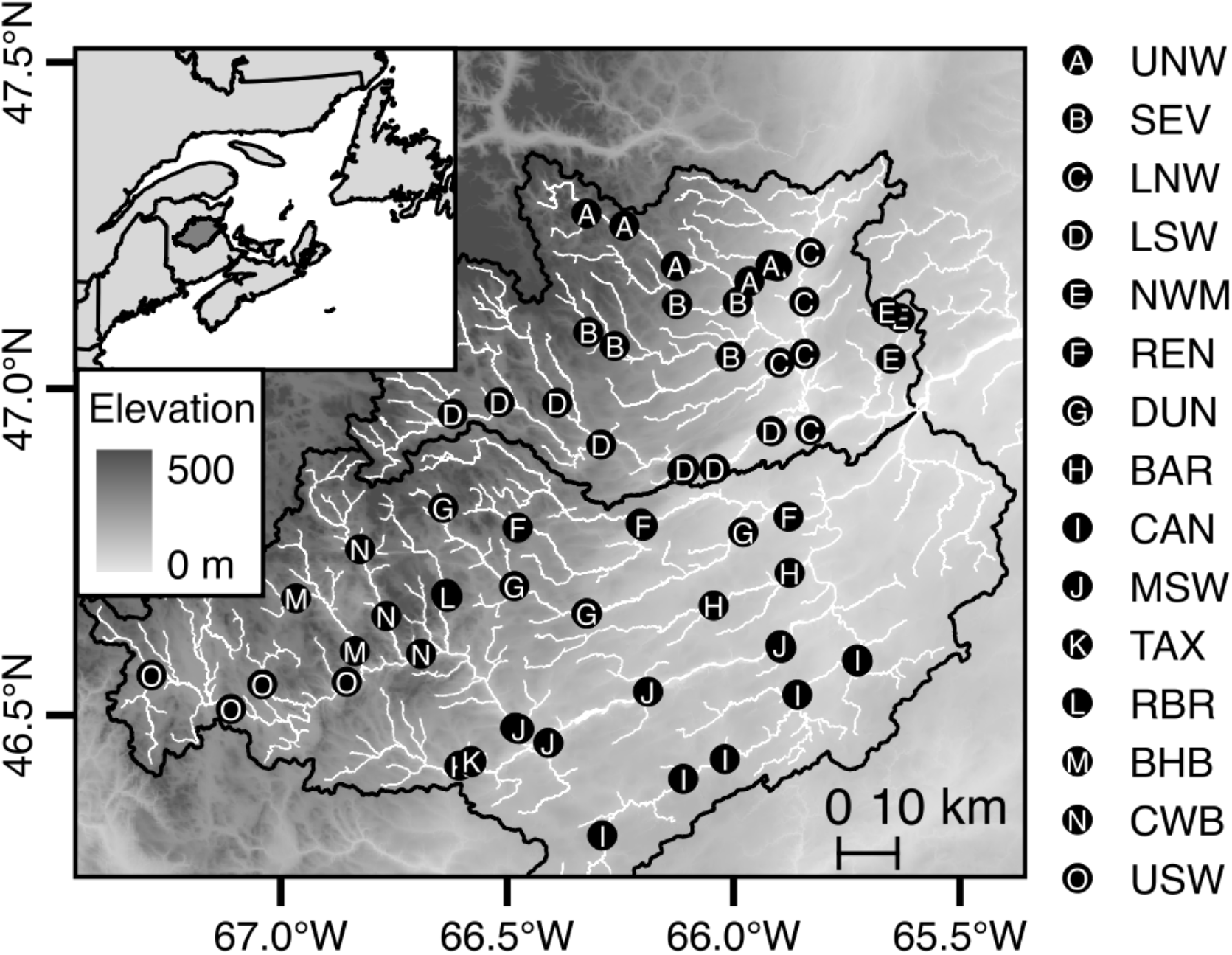
Map of sampling locations for Atlantic salmon and tributary groupings in the Miramichi River (see Table 1 for description of tributary codes). Watersheds of the Northwest and Southwest branches of the river are outlined in black and shading indicates elevation above sea-level.

To simplify interpretation of population structure in this system, we grouped sampling sites together to represent 15 major tributaries within the Miramichi River system. While our choice of the hierarchical level at which to group samples is necessarily arbitrary, our grouping of sampling sites according to major watersheds generally reflects the decisions that returning adults would need to make as they migrate to spawning areas and thus the potential for development of within-river population structure. Some of these groupings occur over long stretches of the same river and may cause genetically distinct groups to be inadvertently clumped together; however, the prevailing view of hierarchical population structure for salmon in river systems suggests that our groupings will have captured a coarser level of population hierarchy if it is present in the system. Where possible we include analyses using site or individual level data to provide confidence these grouping do not influence the patterns of genomic variation we observe.

DNA was extracted from fin clips by the Center for Aquaculture Technologies Canada (Souris, PEI) using a standard DNA extraction procedure and samples were genotyped using a custom 50K SNP Axiom genotyping array designed for North American Atlantic Salmon at the Centre for Integrative Genetics (CIGENE, Norwegian University of Life Sciences, Norway). Raw data from the arrays was processed following the Affymetrix Best Practices Workflow using thresholds of >0.8 Dish Quality Control (DQC) and >0.97 Quality Control (QC) call rate for high-quality samples. A number of samples (163/766) failed to meet these thresholds and thus we attempted to recover data from these sample by re-implementing the Affymetrix Best Practices Workflow with thresholds of >0.8 DQC and >0.90 QC call rate. Samples not exceeding these reduced thresholds (23 samples) were excluded from further analysis. We then compared the minor allele frequency (MAF) of each SNP in all retained individuals under the reduced quality thresholds with the frequency estimated for the high-quality samples and high-quality thresholds. Any SNPs demonstrating significant shifts in allele frequency between the two quality thresholds were imputed as missing genotypes for the recovered samples. SNPs were ordered following their genomic position in the Atlantic salmon reference genome (ICSASG v2; NCBI Genbank accession: GCA_000233375.4; Lien et al., 2016). Due to the commercial nature of the SNP array, SNP positions in the dataset have been shifted randomly by up to 100 bp from their exact location on the reference genome to protect intellectual property while maintaining genomic order and approximate genomic location. We removed both samples and SNPs with greater than 10% missing data as well as SNPs with a MAF less than 0.01 from the data set using PLINK v1.9 (Chang et al., 2015). To remove closely related individuals from the dataset, we implemented an identity-by-descent analysis in PLINK and removed one individual from each pair of individuals with a relatedness coefficient greater than 0.45.

For population genetic analyses that assume a set of neutral and unlinked markers, we created a reduced dataset. We first used BayeScan v2.1 (Foll & Gaggiotti, 2008) with program defaults (20 pilot runs of 5,000 iterations followed by 50,000 burn-in iterations and 50,000 iterations sampled every 10 iterations) but with relaxed settings (prior odds = 10) to liberally characterize F_ST_ outlier markers globally among all the tributary samples. Convergence of the MCMC chain was assessed and confirmed by visually inspecting the posterior parameter distributions. These markers were first removed from the dataset and we then removed SNPs with high physical linkage using a sliding-window approach in PLINK where SNPs with a variance inflation factor greater than two (VIF > 2) were removed from 50 SNP windows shifted by 5 SNPs after each iteration. We further removed all SNPs on the fused chromosomes 8 and 29 (see Methods and Results below). This neutral and LD pruned dataset is hereafter referred to as the neutral dataset and, unless otherwise stated, was used as the primary dataset for population genetic analyses.

### Detection and characterization of structural variation

We used principal components analysis (PCA) implemented in PLINK to visually explore patterns of allelic variation for the whole SNP dataset. The first two PCs of a PCA on the entire dataset revealed a striking pattern of three groups of samples that could not be explained by biological characteristics (e.g. weight, length, sex) or geographic grouping of the samples (see Results). We investigated this pattern further, first using pcadapt v3.0.4 (Luu, Bazin, & Blum, 2017) to characterize divergence between these three groups. Visual inspection of PC axes 3+ indicated that only axes 1 and 2 were involved in generating the three groups and thus we set the number of PCs retained for pcadapt to K = 2. This analysis revealed two strongly divergent genomic regions localized to the beginnings of chromosomes 8 and 29 (F_ST_ > 0.6). An aquaculture population of Atlantic salmon with North American origins have previously been characterized as polymorphic for a fusion between chromosomes 8 and 29 (Brenna-Hansen et al., 2012). Thus, we further investigated to see if variation in chromosome structure could explain the observed pattern in our data. We first calculated pairwise linkage (r^2^) among all SNPs located on chromosomes 8 and 29 using PLINK. Based on the pattern observed, we reversed the order of chromosome 8 and digitally merged it with chromosome 29. We then used the package inveRsion v1.22 (Cáceres, Sindi, Raphael, Cáceres, & González, 2012) in R v3.3.3 (R Core Team 2017) to estimate the break points delineating a chromosomal rearrangement (block size = 5 SNPs, window = 2 Mb, BIC threshold = 100). Using the identified break points, we then assigned a rearrangement genotype (karyotype) to each individual using the R package invClust v1.0 (Cáceres & González, 2015). We used karyotype as a grouping factor and investigated patterns of genetic diversity across chromosomes 8 and 29. Minor allele frequency and heterozygosity for each karyotype, as well as pairwise F_ST_ among karyotypes were calculated using PLINK.

Based on individual karyotype assignments, we calculated the frequency of the fusion (called allele “A”) and the relative proportion of the three karyotypes (AA: homokaryote fused, AB: heterokaryote, BB: homokaryote unfused) for each tributary. Heterogeneity in fusion frequencies between tributaries was tested using an analysis of deviance on a generalized linear model (GLM) with binomial logistic transformation. Within each tributary, we tested for deviation from Hardy-Weinberg equilibrium (HWE) expectations using a chi-square test. We also tested for population structure among tributaries based on frequencies of fusion karyotypes as detailed below for neutral markers. North American Atlantic salmon are also known to be polymorphic for both a chromosomal fusion between chromosomes 26 and 28 as well as a translocation of the p-arm from chromosome 1 to chromosome 23 (Brenna-Hansen et al., 2012). We investigated patterns of linkage between these sets of chromosomes as described above but found no similar patterns as those described for chromosomes 8 and 29.

### Tributary genetic diversity and population structure

Observed and expected heterozygosities as well as inbreeding coefficients were calculated for each of these populations using PLINK. The effective number of breeders (Nb) for the sampled cohort was estimated using the bias-corrected linkage disequilibrium method of Waples and Do (2010) assuming random mating with an allele frequency threshold of >= 0.05, as implemented in the software NeEstimator v2.1 (Do et al., 2014). Genomic datasets with thousands of markers are known to introduce a downward bias for Nb estimates due to linkage disequilibrium caused by the inclusion of comparisons of physically linked markers (Waples, Larson, & Waples, 2016). Despite our creation of a neutral dataset with low physical linkage, we used genomic position information of the SNPs to only compare linkage among SNPs located on different chromosomes and thus absolutely not in physical linkage. Due to the evidence for a fusion between chromosomes 8 and 29, we further excluded these chromosomes from the dataset used to calculate Nb. We then corrected Nb to account for overlapping generations in the parents of the sampled cohort following Waples et al. (2014) and using parameters derived by Ferchaud et al. (2016) for Atlantic salmon.

We calculated Weir and Cockerham’s (1984) unbiased estimate of F_ST_ among all pairs of tributary populations and the statistical significance of F_ST_ estimates was assessed using 10,000 permutations of the data in Arlequin v3.5.2.2 (Excoffier & Lischer, 2010). We applied a FDR correction to account for multiple tests and assess statistical significance of pairwise F_ST_ values (Benjamini & Hochberg, 1995). To explicitly test for hierarchical population structure, we used an AMOVA where tributary populations were grouped according to their location in the two main branches of the river (Northwest and Southwest). The tributary NWM was excluded from this analysis because it is the recipient stream for residual stocking product (i.e. it receives hatchery-derived salmon from mixed and untraceable stocks). AMOVA calculations were conducted in Arlequin v3.5.2.2 and 10,000 permutations were used to assess the statistical significance of population structure.

To test for a pattern of isolation by distance among tributary populations, we used a Mantel test as well as the Redundancy Analysis (RDA) approach recommended by Meirmans (2015). We used the riverdist v0.15.0 (Tyers 2017) package in R v3.3.3 (R Core Team 2017) to calculate the shortest linear river distances between tributary sampling sites. We tested the association of pairwise linear river distance with linearized pairwise genetic distance (F_ST_/1-F_ST_) using a Mantel test with 1,000 permutations as implemented in the R package vegan v2.4-5 (Oksanen et al. 2017). We also decomposed the genetic variance explained by geographic distance using RDA as implemented in vegan v2.4-5. We first calculated distance-based Moran’s eigenvector maps (MEMs) from the linear river distances among sites using the R package adespatial v0.1-0 (Dray et al. 2017) to represent the sampling sites in Cartesian coordinates that accounted for dispersal restrictions induced by the constraints of the dendritic structure of the river. The MEMs with a positive value of Moran’s I were then used as predictor variables in an RDA where the tributary minor allele frequencies were used as the dependent variables. We used a forward step procedure to only select MEMs that explained significant (1,000 permutations, p < 0.05) variance in allele frequencies. Significance of the final model, constrained axes, and the marginal significance of selected MEMs were assessed using 1,000 permutations. The proportion of among-population genetic variation explained by river distance is thus the proportion of total RDA variance that is explained by constrained axes in the final model.

### Clustering analyses

We used several naïve clustering routines to also test for population structure. We first used Admixture v1.3 (Alexander, Novembre, & Lange, 2009) to estimate the number of genetic clusters in the neutral dataset. Admixture was with 5-fold cross-validation for a range of putative populations (K = 1 to K = 10). The number of genetic groups (K) was chosen based on the lowest cross-validation error. We also used Discriminant Analysis of Principal Components (DAPC; Jombart, Devillard, & Balloux, 2010) to provide a non-model-based approach for estimating genetic clusters. The number of genetic clusters was determined using k-means clustering of principal components transformed allele frequency data, as implemented in the ‘find.clusters’ function of the adegenet v2.1.0 package (Jombart & Ahmed, 2011) in R v3.3.3 (R Core Team 2017). The number of clusters was determined based on the profile of the Bayesian information criterion (BIC) across a range of putative populations (K = 1 to K = 10). Finally, we used principal components analyses (PCA), as implemented in PLINK, to visually investigate population groupings in multivariate allelic space. To assess the robustness of our PCA analysis, we also used the ‘smartpca’ algorithm implemented in EIGENSOFT (Patterson, Price, & Reich, 2006) which prunes outlier samples and SNPs and we conducted PCAs using MAF thresholds of 5%, 10% as well as on the full dataset.

### Outlier analyses

The restricted dispersal of organisms in river networks results in inflated variance in F_ST_ at neutral markers compared to that expected under traditional population genetic models and leads to increased false positive outliers if these demographic relationships are not properly take into account (Fourcade, Chaput-Bardy, Secondi, Fleurant, & Lemaire, 2013). To reduce the detection of false positive outliers we implemented two outlier tests that take into account the genetic relatedness of populations: the modified Lewontin-Krakauer (FLK) test of Bonhomme et al. (2010) and BayPass v2.1 (Gautier, 2015). These test control for demography by taking into account the genetic relatedness of populations and thus should exhibit reduced false-positives relative to tests assuming an island model (e.g. BayeScan). For FLK, we first estimated Reynold’s distance among populations based on the neutral SNP dataset (20,967 SNPs) and then calculated the T_F-LK_ statistic for each SNP as implemented in R following Bonhomme et al. (2010). R code for the FLK analysis was obtained from: https://qgsp.jouy.inra.fr (date accessed: December 15, 2017). Statistical significance of outliers was assessed by comparing the test statistic to the Chi-squared distribution and correcting the p-values using FDR of 0.05. BayPass was run using program defaults (20 pilot runs of 1,000 iterations each to optimize MCMC parameters proposal distributions followed by a 5,000-iteration burn-in and finally 25,000 iterations sampled every 25 steps). We ran the program five times with different starting seeds and confirmed convergence of the algorithm by assessing correlations of the estimated parameters among runs. BayPass does not provide p-values and thus we used the ranking of loci based on the divergence statistic (XtX) to corroborate results from other tests. Due the lack of hierarchical structure we also conducted outliers tests using BayeScan v2.1 (Foll & Gaggiotti, 2008) with program defaults (20 pilot runs of 5,000 iterations followed by 50,000 burn-in iterations and 50,000 iterations sampled every 10 iterations) but with the prior odds set to 10,000 to reduce the incidence of false positives (Lotterhos & Whitlock, 2014). We assessed and confirmed convergence of the MCMC by visually inspecting the posterior parameter distributions. OutFLANK was also used to detect outlier SNPs (Whitlock & Lotterhos, 2015). Loci putatively experiencing divergent selection were identified as those with FDR < 0.05. We identified a consensus set of outliers as the intersection of significant outliers across all outlier approaches. To identify genes that are potential targets of selection we used BEDTools v2.26.0 (Quinlan & Hall, 2010) to search the Atlantic salmon genome for genes that occurred within a 10 Kb window on either side of each outlier.

### Multivariate selection and environmental association

Based on the established roles of climate and stream conditions in promoting local adaptation of Atlantic salmon populations (Bourret, Dionne, et al., 2013; Garcia de Leaniz et al., 2007; Jeffery et al., 2017), we also investigated spatial patterns of association between genomic variation and climatic (i.e. temperature and precipitation) and physiographic (i.e. river distance and elevation above sea level) attributes of our sampling sites. We employed a polygenic framework based on Redundancy Analysis (RDA) to test for signatures of multivariate selection (Forester, Lasky, Wagner, & Urban, 2018). We also tested for correlations of the climate and physiography variables with the frequency of the fusion and outlier SNPs allele frequencies using logistic regression.

First, to characterize climatic and physiographic attributes of our sampling sites we used GPS locations of the sampling sites to extract temperature and precipitation normals (1970 – 2000) from the WorldClim 2.0 Bioclimatic database (Fick & Hijmans, 2017), as well as elevation (NRCAN 2016) using QGIS v2.14 software (QGIS Development Team 2016). We calculated distance to the river mouth using the riverdist v0.15.0 package in R. Because tributaries are represented by more than one sampling site and our allele frequency data was calculated on the basis of tributaries, we summarized the environmental data for each tributary by taking a weighted average of the sampling site level environmental data weighted by the number of individuals at a site for each tributary. While this may result in a loss of power by summarizing variation over larger spatial areas our weighted averages should reflect the composite effects of selection on alleles across sampling sites and provide a conservative estimate of SNP-environment associations. Many of the climate and physiography variables were highly correlated across tributaries (|r| > 0.8; Figure S1), so we first performed a variable reduction step by using two PCAs to summarize environmental variation for combined temperature/elevation/distance variables and the precipitation variables separately. For each of these PCA, we retained PC axes based on the broken-stick distribution (Frontier, 1976). Site scores of the retained PC axes represent summaries of environmental variation and were subsequently used as predictors of genomic variation.

Multivariate polygenic selection was tested using RDA with three environmental summary variables (see results) as predictors and tributary allele frequencies for the whole SNP dataset as dependent variables. We performed forward step selection of the environmental variables and retained those that explained significant (1,000 permutations, p < 0.05) variance in allele frequencies. Variance inflation factors were below 10 for all variables indicating no multicollinearity between the predictors. Significance of the final model was assessed using 1,000 permutations. To identify SNPs influenced by environmental variation we scaled and centered the raw scores for SNPs on each constrained RDA axis. We then used a robust Mahalanobis distance metric similar to that applied in the outlier software pcadapt (Luu et al., 2017) to identify SNPs significantly loaded on each constrained axis. Complex spatial environments like rivers can generate population structure that interferes with the ability to detect environmental associations; however, Brauer et al. (2018) have demonstrated the utility of partial RDA to increase resolution of genotype-environment associations in stream networks. To control for population structure while testing for environmental associations, we performed a partial RDA as described above while using the significant MEMs from the isolation by distance analysis as conditioning variables as suggested by Brauer et al. (2018).

To test the individual association between the chromosome 8 and 29 fusion or outlier SNP frequencies and environment, we used a GLM with a logistic link function for binomial data, the response variable being the number of individuals carrying/not carrying the fusion arrangement or the alternative allele and the explanatory variable being an environmental variable. Given that outlier SNPs on the same chromosome were in strong linkage disequilibrium and that the distribution of frequencies was highly correlated (Figure S2), we performed the analysis on the mean MAF across outlier SNPs from the same chromosome. We corrected p-values for multiple comparisons following Benjamini and Hochberg (1995).

### Functional enrichment

To test for functional enrichment of genes associated with reduced recombination region of the fusion as well as the significant environmental variables from the RDA we conducted Gene Ontology enrichment tests. We first extracted gene ontology (GO) information for all Atlantic salmon genes from SalmoBase (Samy et al., 2017; www.salmobase.org, accessed: November 25, 2017). We then determined reference sets of genes for each analysis as all of the Atlantic salmon genes for the fusion and all genes within 10 Kb of a SNP on our array for the environmental outliers (the same window used to associate genes with significant SNPs). We tested for functional enrichment of GO biological process terms using the “weight01” algorithm to account for the structure of the GO term relationships as implemented in the topGO v2.26 package in R (Alexa, Rahnenführer, & Lengauer, 2006).

## Results

### Sampling and SNP genotyping

A total of 766 age 1+ Atlantic salmon were collected throughout the Miramichi River watershed. The distributions of length, weight, and condition factor were unimodal for most tributaries and indicate the collections primarily represented a single cohort (overall mean ± sd; fork length = 85.1 ± 6.7 mm; weight = 7.4 ± 1.7 g; Figures S3, S4, S5). Only fork length in the Renous River sample (REN) deviated from unimodality (p = 0.04; all other tributaries p > 0.14), but this difference was not significant following FDR correction. While we did not directly age the fish in our collections, there is a well-established length-age relationship for fish in this river system that has previously been validated by scale aging (Swansburg et al., 2002). The lengths of our 1+juveniles are consistent with the size distribution of 1+juveniles from the Miramichi River over a 30 year period and fall well within the interannual variation for these measures (Figure S3; Swansburg et al., 2002). While there is some known overlap in the size distribution of 1+ and 2+ juveniles (Swansburg et al., 2002), we only collected 32 individuals with a fork length larger than 95 mm suggesting if some of these fish do represent 2+ juveniles, the overall contribution of 2+ juveniles to our collections are limited.

Size of fish varied among tributaries (Fork Length: F14,751 = 5.7, p < 0.001; weight: F14,751 = 5.4, p < 0.001; condition factor: F14,751 = 12.3, p < 0.001; Figures S3-S5). These patterns were driven by slightly smaller lengths and weights of fish from Rocky Brook (RBR; Figures S3, S4); however, condition factor differences were driven by the generally higher condition for fish from the Upper Northwest Miramichi River (UNW; Figure S5). The relative proportions of males to females was similar for most tributaries (*χ*^2^ = 21.8, p = 0.08); however, the proportion of precociously mature males varied considerably among tributaries (*χ*^2^ = 52.9, p < 0.001; Figure S6). Controlling for sex and precocious maturation status did not have an effect on the differences in size observed among tributaries (results not shown).

There were 23 samples we were unable to genotype due to the genotyping arrays not meeting the quality standards described in the methods and a further 15 individuals were removed from the dataset for having more than 10% missing data. The identity-by-descent analysis identified 41 potential full-sib pairs and removal of 26 individuals eliminated these relationships. We retained data for 702 individuals at 52,537 SNPs in the full dataset. The overall genotyping rate was high with less than 2% missing data in the full dataset. Using nonconservative settings to detect putative outlier (likely with many false positives) we identified and removed 259 outliers. Following position-based pruning for linkage disequilibrium (N = 30,080) and removal of SNPs (N = 1483) on the fused chromosomes 8 and 29, a total of 20,974 SNPs were retained in the neutral dataset that was used for most population genetic inferences.

### Detection and characterization of structural variation

Exploratory analyses of the entire dataset using PCA revealed three prominent clusters that did not obviously reflect geographic (i.e. tributaries) or biological characteristics (i.e. sex, length, weight) of the samples (Figure 2A). Divergence among these three groups was driven entirely by two regions located on chromosomes 8 and 29 (Figure 2B). Mean divergence between these three groups within this region (F_ST_ = 0.27) was nearly three orders of magnitude greater than mean divergence between these groups across the rest of the genome (F_ST_ = 0.0003). Pairwise LD among SNPs on chromosomes 8 and 29 revealed two prominent linkage blocks, each spanning several Mb, within both chromosomes and strong linkage (mean pairwise r^2^ within the region = 0.17 vs. mean pairwise r^2^ outside the region = 0.007) between 8 Mb of chromosome 8 and 3 Mb of chromosome 29 (Figure 2D). This pattern reflects the fusion between chromosomes 8 and 29 that has been previously described in a North American Atlantic salmon aquaculture population using linkage mapping and cytogenetic techniques (Brenna-Hansen et al., 2012). Our patterns of linkage suggest that the p-arm of chromosome 8, containing primarily rDNA repeated units (Lien et al., 2016; Pendás, Morán, & Garcia-Vázquez, 1993), fused with the centromere of the acrocentric chromosome 29 (Figure 2D).

**Figure 2:**
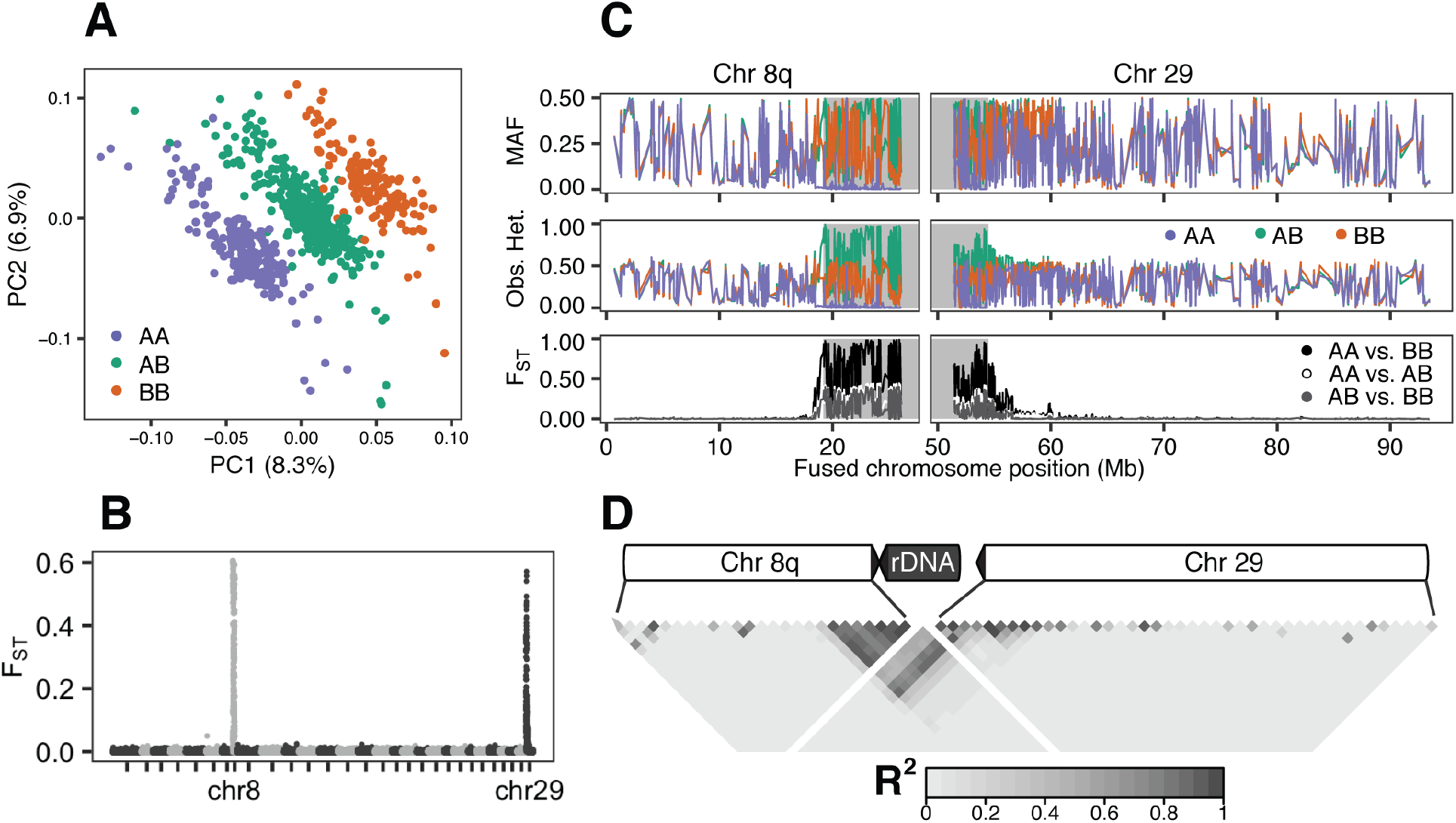
Characterization of a fusion between Atlantic salmon chromosomes 8 and 29. (A) Principal components analysis revealed three prominent groups corresponding to homokaryote fused (purple / top right), homokaryote unfused (green / bottom left), and heterokaryote (orange / middle) individuals. (B) Genetic differentiation (FST) between the three karyotypes is localized on chromosomes 8 and 29. (C) Genetic diversity (minor allele frequency = MAF, observed heterozygosity = Obs. Het.) is reduced for the fused homokaryotes (AA, purple) and genetic divergence (FST) between homokaryotes (AA vs. BB, green) is elevated in a region of high linkage among chromosomes (grey shading). (D) Linkage disequilibrium (R^2^) summarized in 1 Mb windows within and between chromosomes 8 and 29. Patterns of linkage suggests the p-arm of chromosome 8 containing rDNA repeated elements fused to the centromere of the acrocentric chromosome 29.

This fusion has not been fixed in North American populations and thus individuals may carry two 8-29 fused chromosomes (AA), two unfused sets of chromosomes 8 and 29 (BB) or one fused 8-29 and one unfused set of chromosomes 8 and 29 (AB). Principal components analysis using only the SNPs contained within the linked regions revealed three distinct groupings allowing unambiguous assignment of individuals to the three karyotypes (Figure S7). Genetic diversity within each of the three karyotype groups revealed that fused homokaryotes have dramatically reduced allelic diversity and heterozygosity in the highly linked region while unfused homokaryotes do not exhibit this pattern (Mean heterozygosity within vs. outside linked region: AA = 0.08 vs. 0.27, t = 22.3, p ≪ 0.001; BB = 0.31 vs. 0.29, t = 1.8, p = 0.11; Figure 2C). These results are consistent with observations that recombination is reduced near the fusion point in fused chromosomes but not in unfused chromosomes (Bidau, Giménez, Palmer, & Searle, 2001; Dumas & Britton-Davidian, 2002).

The fusion karyotypes were in Hardy-Weinberg equilibrium in all tributary populations and there was no sex bias in the distribution of fusion (*χ*^2^ = 0.35, df = 2, p = 0.84). The fusion was polymorphic in all tributary samples and its frequency was significantly heterogeneous between tributaries (deviance = 39, df = 14, p < 0.001), ranging from 38% to 66%. However, there was no overall difference in fusion frequency between the two main branches of the river (NW vs. SW, *χ*^2^ = 1.95, df = 2, p = 0.38). Further population structure and environmental associations with the fusion karyotype are presented below with the results for SNPs.

### Tributary genetic diversity and population structure

The tributary samples exhibited similar levels of observed and expected heterozygosity and levels of inbreeding were low (Table 1). The effective number of breeders ranged over two orders of magnitude from approximately 80 in Taxis River to over 4000 in Clearwater Brook (Table 1). Genetic divergence among tributaries was low, but statistically significant (Global F_ST_ = 0.004, pairwise F_ST_ range = 0.001 – 0.014; Figure 3). Three tributary populations from upstream reaches exhibited consistent stronger differentiation from all other tributaries, i.e., in the Southwest Miramichi: Rocky Brook (RBR, mean F_ST_ = 0.01), and Taxis River (TAX, mean F_ST_ = 0.007) and in the Northwest Miramichi: Upper Northwest Miramichi River (UNW, mean F_ST_ = 0.006; Figure 3). Additionally, Northwest Millstream which empties into the tidal reaches of the Northwest Miramichi and regularly receives hatchery residuals of untraceable origin exhibited similar levels of differentiation (NWM, F_ST_ = 0.007). Omitting these populations reduced the magnitude of overall structure by half (F_ST_ = 0.002). Patterns of pairwise F_ST_ are similar when conducted at the sampling site level for sites with more than 12 individuals indicating our grouping of sites is not driving the patterns of structure we report (Figure S8). Hierarchical structure was not supported by the AMOVA as no genetic variance was explained by the two main branches of the Miramichi River (F_ST_ = 0.0001, p = 0.35; Table 2); however, the AMOVA did support population structure among tributaries within the main branches of the river consistent with the pairwise F_ST_ results (F_ST_ = 0.004, p < 0.001; Table 2). The Mantel test for isolation by river distance was non-significant (p = 0.17, Mantel’s r = 0.09); however, the spatial RDA identified the first MEM as explaining significant among-population variation in allele frequencies (p = 0.03; Figure 4). The spatial RDA resulted in a single constrained axis that explained 1.1% of the among-population variation in allele frequencies. This weak pattern of spatial structure was driven by differentiation of the tributaries in the upstream reaches of the Southwest from the downstream Southwest tributaries and Northwest tributaries (Figure 4). To ensure robustness of the analyses conducted on the neutral dataset and that our filtering thresholds did not remove important signals of population structure we also performed these analyses on both the full dataset and a dataset with less stringent linkage filtering (VIF > 2 in 10 Kb windows). These analyses resulted in nearly identical results for all analyses (results not shown) and indicate the patterns reported here are robust.

**Figure 3:**
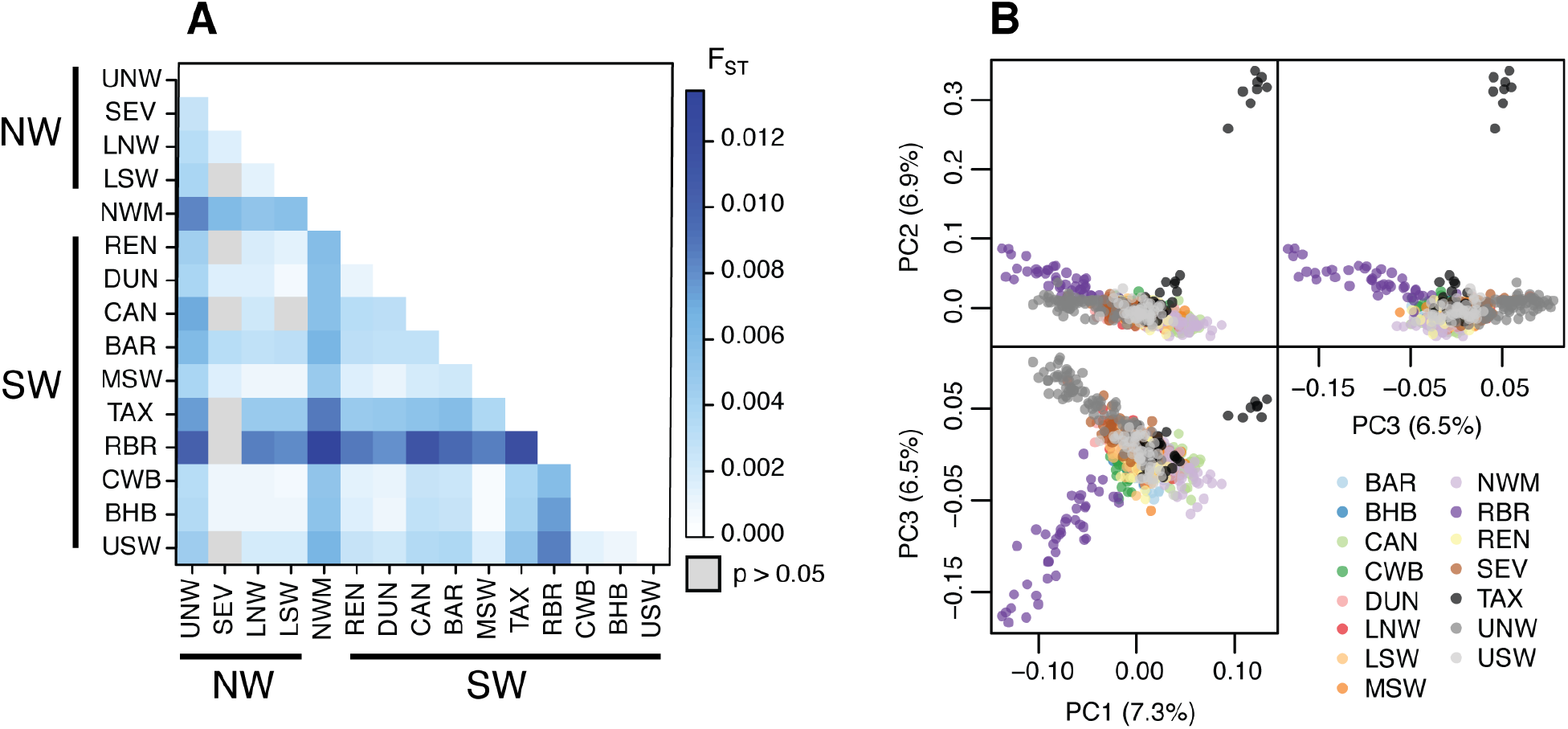
Genetic structure of Atlantic salmon in the Miramichi River based on neutral LD-pruned SNPs. (A) Pairwise F_ST_ among 15 tributary populations. Populations arranged in geographical order from upstream in the northwest branch (NW) to upstream in the southwest branch (SW). See Table 1 for description of site codes. Grey boxes indicate F_ST_ comparisons that are not statistically different from zero based on 10000 permutations of the data. (B) Principal components analysis of allelic variation. Color indicates tributary of sampling, see Table 1 for description of site codes.

**Figure 4:**
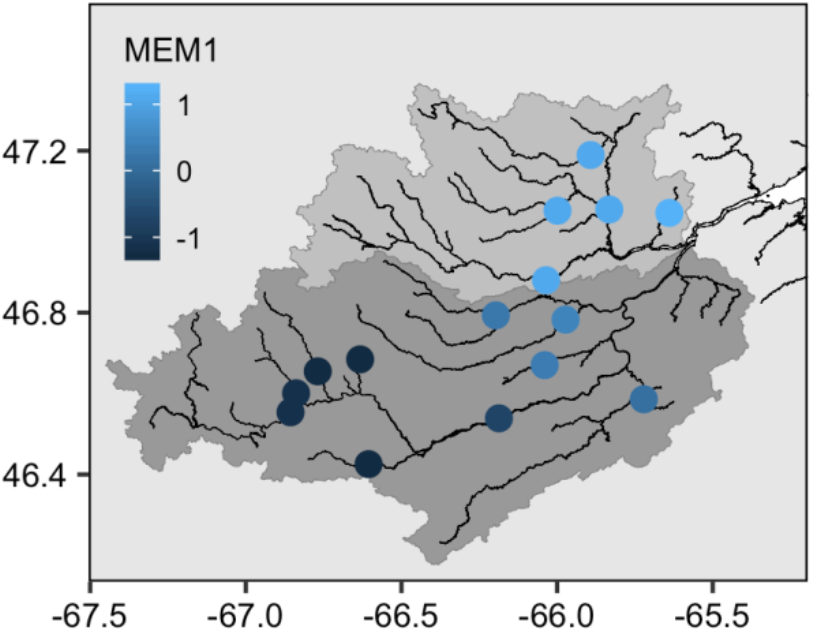
Weak spatial structuring of neutral genetic diversity. The first Moran’s eigenvector map (MEM1) of pairwise river distance among tributaries explains 1.1% (RDA, 1000 permutations, p = 0.03) of the allelic variation among tributaries. Site loadings on MEM1 (color scale) generally separate tributaries in the upper reaches of the Southwest Miramichi River (dark grey shading) from tributaries in the lower Southwest and Northwest Miramichi River (medium grey shading).

**Table 1:** Genetic diversity summary statistics for Atlantic salmon populations from 15 tributaries of the Miramichi River. Populations ordered geographically from North to South and grouped according to the two main branches of the river. Northwest Millstream enters into the estuary and thus was not included in either branch (see Figure 1).

| Tributary | Code | N | H<sub>O</sub> | H<sub>E</sub> | F<sub>IS</sub> | N<sub>b</sub><sup>Cor</sup> |
| --- | --- | --- | --- | --- | --- | --- |
| Northwest Millstream | NWM | 43 | 0.297 | 0.280 | -0.07 | 145 |
| *Northwest branch* |  |  |  |  |  |  |
| Upper Northwest | UNW | 77 | 0.287 | 0.283 | -0.02 | 723 |
| Sevogle | SEV | 48 | 0.289 | 0.285 | -0.03 | 635 |
| Lower Northwest | LNW | 62 | 0.290 | 0.286 | -0.02 | 630 |
| Little Southwest | LSW | 133 | 0.294 | 0.286 | -0.02 | 494 |
| *Southwest branch* |  |  |  |  |  |  |
| Renous | REN | 48 | 0.291 | 0.284 | -0.03 | 440 |
| Dungarvon | DUN | 49 | 0.290 | 0.285 | -0.03 | 1783 |
| Bartholomew | BAR | 25 | 0.293 | 0.284 | -0.07 | 556 |
| Cains | CAN | 55 | 0.294 | 0.287 | -0.03 | 678 |
| Southwest | MSW | 70 | 0.295 | 0.287 | -0.03 | 531 |
| Taxis | TAX | 39 | 0.297 | 0.282 | -0.07 | 77 |
| Rocky Brook | RBR | 43 | 0.290 | 0.282 | -0.05 | 108 |
| Clearwater | CWB | 61 | 0.293 | 0.285 | -0.03 | 4300 |
| Burnthill Brook | BHB | 24 | 0.294 | 0.281 | -0.07 | 81 |
| Upper Southwest | USW | 47 | 0.290 | 0.284 | -0.03 | 168 |

N = number of individuals, H<sub>O</sub> = observed heterozygosity, H<sub>E</sub> = expected heterozygosity, F<sub>IS</sub> = inbreeding coefficient, N<sub>b</sub><sup>Cor</sup> = corrected effective number of breeders.

The population structure among tributaries based on the frequency of the chromosomal fusion was five times stronger than that exhibited by SNPs (global F_ST_ = 0.019 vs. 0.004) and was in the upper 5% of the distribution of global F_ST_ estimates for all SNPs (Figure S9). The estimates of pairwise tributary divergence were not correlated between the fusion and SNP markers (Mantel’s r = 0.19; p = 0.16). AMOVA based on the fusion karyotype variation also failed to support the hierarchical structuring between the main branches (NW vs. SW; F_ST_ = −0.0004, p = 0.66; Table 2) of the Miramichi River but did identify structuring among tributaries within the main branches (F_ST_ = 0.019, p < 0.001; Table 2). The fusion karyotype also did not exhibit a pattern of isolation by distance (Mantel’s r = −0.09, p = 0.83).

**Table 2:** Analysis of molecular variance (AMOVA) of tributary populations in the Miramichi River based on neutral SNPs and chromosome 8 and 29 fusion karyotype.

| Source of variation | % Variation | F-statistic | P-value |
| --- | --- | --- | --- |
| *Neutral SNPs* |  |  |  |
| Among main river branches | 0.01 | 0.0001 | 0.35 |
| Among tributaries within river branches | 0.42 | 0.0042 | <0.001 |
| Among individuals within tributaries | 99.57 | - | - |
| *Chromosome 8 & 29 fusion* |  |  |  |
| Among main river branches | -0.40 | -0.004 | 0.66 |
| Among tributaries within river branches | 1.90 | 0.019 | <0.001 |
| Among individuals within tributaries | 98.50 | - | - |

### Clustering analyses

Principal components analysis of the putatively neutral dataset revealed groupings of samples in multivariate allelic space that reflected the geographic location of sampling (Figure 3B). The first three PC axes explained 7.3%, 6.9%, and 6.5% of the total variation, respectively, and were sufficient to visually separate the Taxis River, Rocky Brook, Upper Northwest Miramichi River, and, to a lesser extent, Northwest Millstream from each other and from the rest of the samples (Figure 3). The PCA results were consistent with the population structure analyses, where most individuals were weakly differentiated but these four tributary populations exhibited elevated divergence relative to the other tributaries. PCAs conducted using EIGENSOFT, inspection of additional axes, and PCAs following removal of divergent populations all failed to reveal additional groupings of individuals (Figure S10). Both more stringent MAF filters (5% and 10%) and no MAF filters used for PCAs produced quantitatively similar results (results not shown). Loadings on the first two PC axes indicate that SNPs driving the differences observed are distributed throughout the genome (Figure S11). Neither Admixture nor DAPC supported genetic clustering of samples likely due to the weak overall structure we observed in this system (Figures S12 – S14). These results reinforce that the overall weak structure we report is not a result of the grouping of sites within tributaries.

### Outlier analyses

We detected 22, 28, and 16 outliers using Bayescan, OutFlank, and FLK, respectively (Figure S15). In total, 14 of these SNPs were identified as statistically significant by all methods and were the top 14 most divergent SNPs according to the XtX statistic of BayPass. Two of the common outliers were found on chromosome 3, one on chromosome 6, and the remaining 11 outliers were clustered in a region of chromosome 9 that spanned approximately 250 kb. We searched the *Salmo salar* genome annotation for genes that fell within 10 kb on either side of each outlier SNP. Thirteen different genes were in proximity to outlier SNPs and could represent putative targets of selection (Table 3). Of particular note, the region on chromosome 9 includes the gene *six6*, which has previously been linked to age- and possibly size-at-maturity in Atlantic salmon in Europe (Barson et al., 2015; Johnston et al., 2014) as well as run-timing (Cauwelier et al., 2017; Pritchard et al., 2018).

**Table 3:**
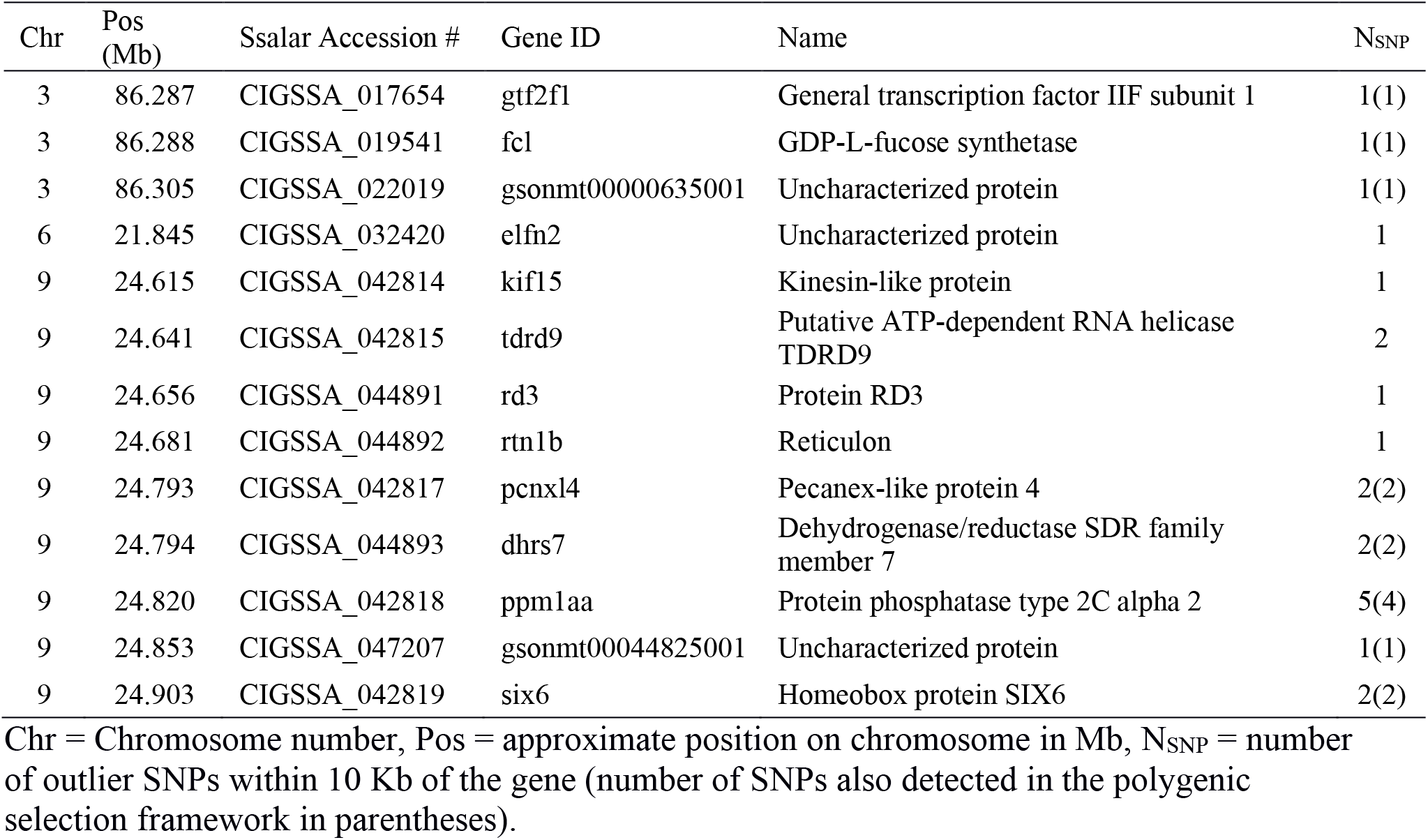
Genes that occur within 10 Kb of common outlier SNPs detected by Bayescan, OutFlank, FLK, and BayPass.

### Multivariate selection and environmental association

The PCA-based reduction of environmental variables resulted in three summary variables (Figure S16) that generally described: 1) temp./elevation, representing the first PC of geographic and temperature data, which explained 73% of variation and was positively associated with temperature (throughout the year) and negatively associated with elevation/distance to river mouth, 2) winter precipitation, PC1 of precipitation data, which explained 52% of variation and was positively associated with precipitation during the coldest/driest months, and 3) summer precipitation, PC2 of precipitation data, which explained 40% of variation and was positively associated with annual precipitation and precipitation during the warmest/wettest months.

When assessing their associations with genomic data, the forward step procedure of the RDA selected only the temp./elevation variable as explaining significant allelic variation and generated one significant discriminant axis (p = 0.001) explaining 3.1% of the total allelic variation. There were 226 SNPs that were significantly associated with variation in temp./elevation. When controlling for underlying population structure using the significant MEM from the spatial RDA, the signal of polygenic selection associated with temp./elevation remained significant (p = 0.014), explaining 2.6% of the allelic variance. After controlling for population structure, variation in temp./elevation correlated with 185 SNPs, including eight outliers (out of 13) previously detected using single locus tests (Table 3; Figure 5).

**Figure 5:**
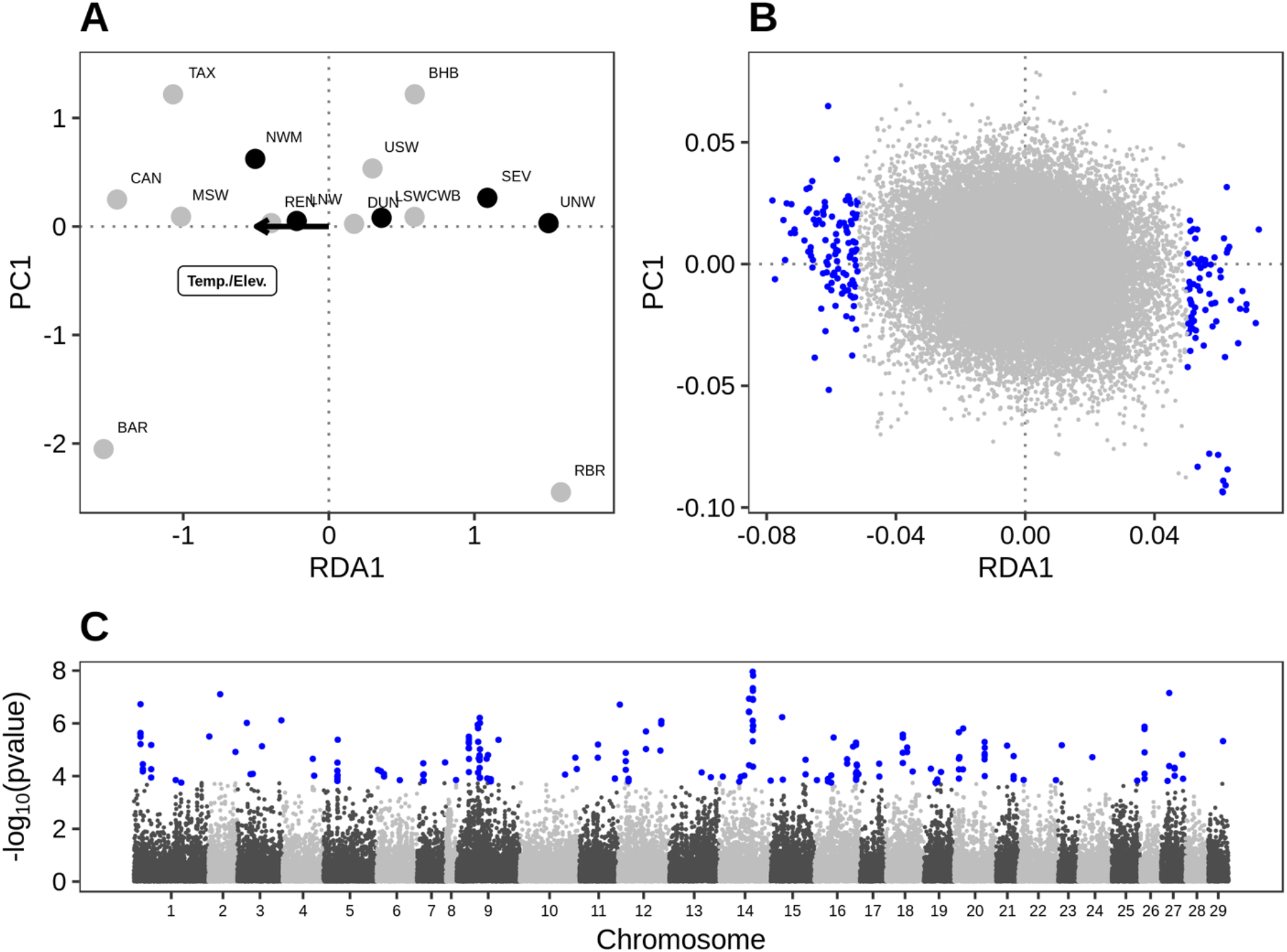
Evidence for polygenic selection among Miramichi River Atlantic salmon populations based on partial redundancy analysis. (A) RDA triplot of the association between tributaries (Northwest = black, Southwest = grey), SNPs, and environmental predictors. The Temp./Elev. summary variable explained 2.6% of the among-population variation in allele frequencies (1000 permutations, p = 0.011). (B) zoom of RDA triplot highlighting 198 outliers (blue points) significantly associated with the Temp./Elev. summary variable (C) Genomic distribution of the strength of SNP association with the Temp./Elev. summary variable (outliers highlighted with blue points).

Consistent with the RDA results, allele frequency variation for outlier SNPs on chromosomes 3 and 6 were significantly associated with variation in temp./elevation (chromosome 3: GLM z = −5.9, p < 0.001, chromosome 6: z = −5.7, p < 0.001, Figure 6A, S17A), which explained 36% (chromosome 3) and 30% (chromosome 6) of allele frequency variation between tributaries in these chromosomal regions. A marginal but significant effect of winter precipitation (GLM z = −2.6, p = 0.009) explains an additional 3-5% of variance for chromosome 6 (Figure 6). On chromosome 9, variation in allele frequency was also significantly associated with temp./elevation (GLM, z = −3.7, p < 0.001) and winter precipitation (GLM, z = 4.3, p < 0.001) and those combined predictors explained 11% of the variation between tributaries on this chromosome.

**Figure 6:**
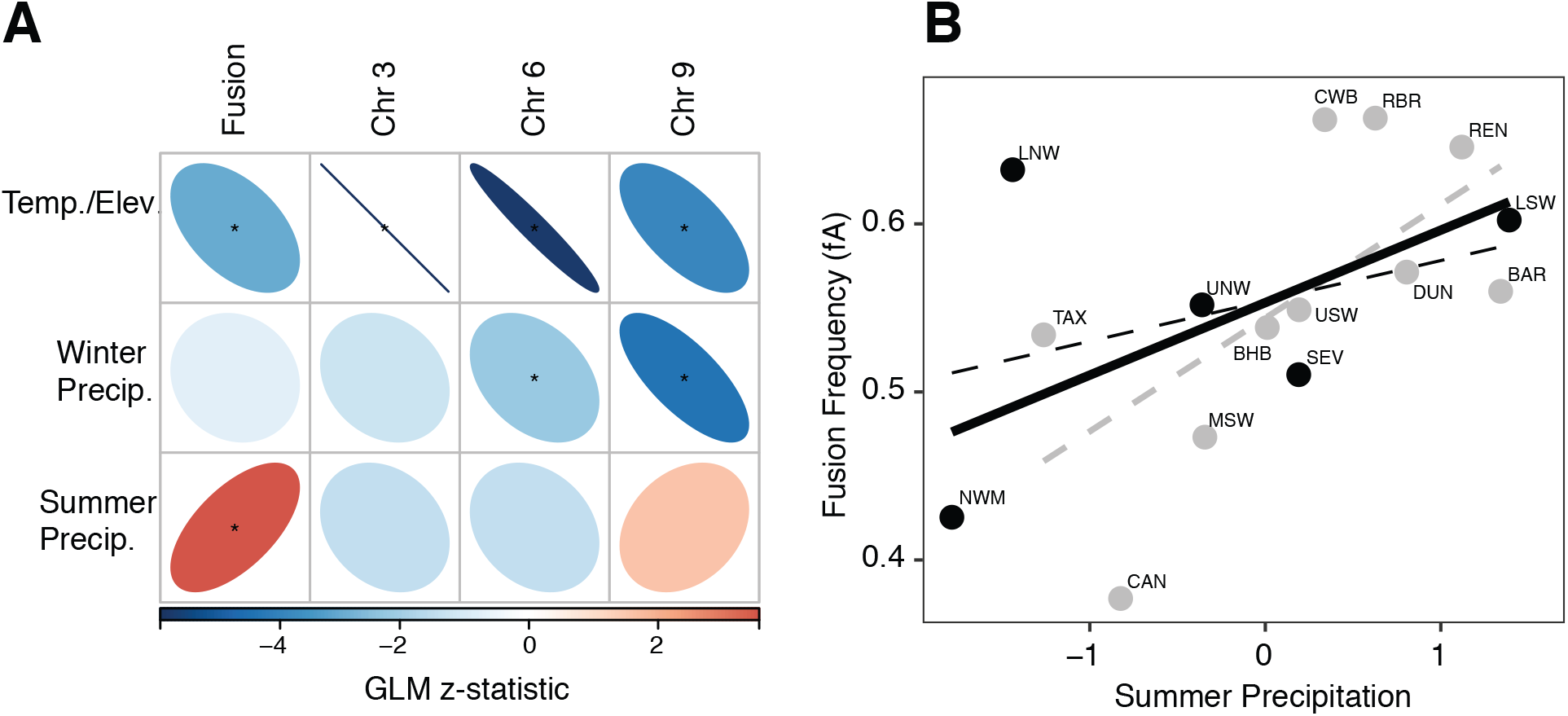
(A) Statistical associations between each environmental predictor and the frequency of fusion rearrangement or the allelic frequency of outlier SNPs. The outlier SNPs on each chromosome were highly linked and thus only one SNP per chromosome is shown (see Figure S11 for specific correlations with each SNP). Strength and direction of the statistical association (GLM) are indicated by the shape of the ellipse and its colour (red: positive, blue: negative). Stars denote significance at 0.05 level. (B) Summer precipitation predicts fusion frequency (solid black line, GLM z = 3.0, p = 0.002, R^2^ = 0.21). This pattern of association is present in both the Southwest branch (grey dashed line, GLM z = 3.6, p = 0.002) and Northwest branch (black dashed line, GLM z = 1.7, p = 0.08) of the river.

Variation in fusion frequency between tributaries was best predicted by variation in summer precipitation, which explained 21% of variance (Figure 6), and by variation in temp./elevation, which explained an additional 5% of variance. The fusion was significantly more frequent in tributaries with higher summer precipitation (GLM, z = 3.0, p = 0.002; Figure 6B) and tributaries at higher elevation characterized by lower temperatures (GLM, z = 2.0, p = 0.02; Figure S17B). Variation in fusion frequencies primarily corresponded to a replacement of one homokaryote by the other homokaryote (Figure S17B). These gradients occurred in parallel for both main branches of the river (Southwest: GLM, z = 3.6, p = 0.002; Northwest: GLM, z = 1.7, p = 0.08; Figure 6B). None of the environmental associations with genetic markers (outlier SNPs or chromosome 8-29 fusion) reflected differences between the NW and SW branches of the river. To verify that these environmental associations were not caused by our choice of sample site groupings we also conducted the analyses at the site level for sites that had a minimum of 12 individuals. In all cases we also detected correlations similar to those we report for tributary-level summaries (e.g. Elevation-six6: GLM, z = 4.0, p < 0.001, Figure S18A; Summer precipitation-fusion: GLM, z = 3.5, p = 0.06; Figure S18B).

### Functional enrichment

We identified 213 genes located within the linked region associated with the chromosomal fusion. These genes represented functional enrichment for 26 GO biological processes (p < 0.01, Table 4). Of particular interest are several processes related to neural development and behaviour, phototransduction, and cell adhesion and cell-cell communication processes involved in contractile tissues. For the 185 genes associated with temp./elevation, 15 GO biological processes were over-represented (p < 0.01, Table 4). Processes related to development and memory dominated the enriched functions.

**Table 4:**
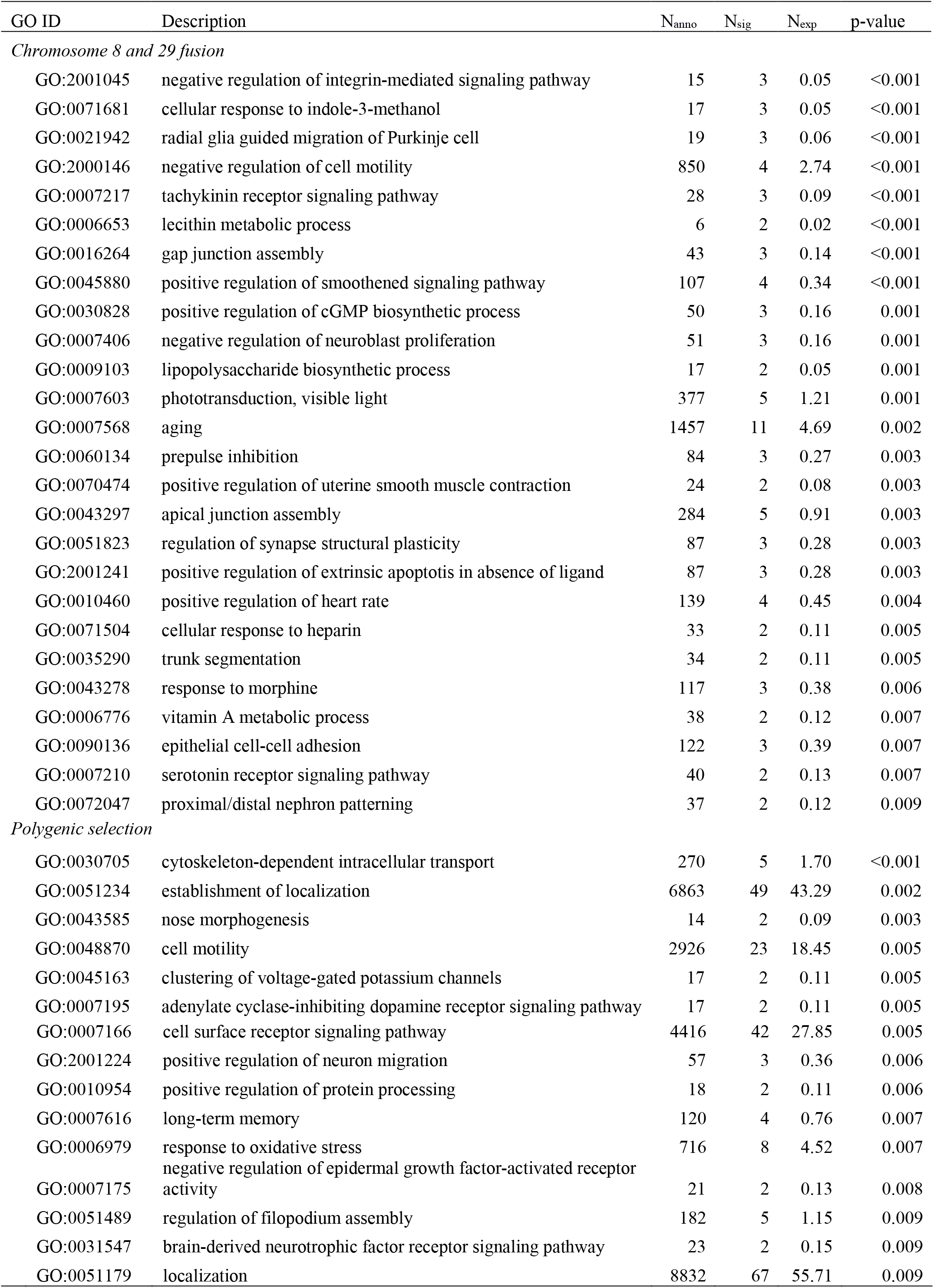
Gene ontology enrichment for genes located within the region of reduced recombination associated with the fusion of chromosomes 8 and 29 and the 198 genes identified as potentially under polygenic selection by their association with an elevation and temperature gradient. N_anno_ = number of genes in reference set annotated with the specific GO function, N_sig_ = number of genes in fusion region with specific GO annotation, and Nexp = number of genes in fusion region with specific GO annotation expected if randomly distributed.

| GO ID | Description | $N_{\text{anno}}$ | $N_{\text{sig}}$ | $N_{\text{exp}}$ | p-value |
| --- | --- | --- | --- | --- | --- |
| *Chromosome 8 and 29 fusion* |  |  |  |  |  |
| GO:2001045 | negative regulation of integrin-mediated signaling pathway | 15 | 3 | 0.05 | <0.001 |
| GO:0071681 | cellular response to indole-3-methanol | 17 | 3 | 0.05 | <0.001 |
| GO:0021942 | radial glia guided migration of Purkinje cell | 19 | 3 | 0.06 | <0.001 |
| GO:2000146 | negative regulation of cell motility | 850 | 4 | 2.74 | <0.001 |
| GO:0007217 | tachykinin receptor signaling pathway | 28 | 3 | 0.09 | <0.001 |
| GO:0006653 | lecithin metabolic process | 6 | 2 | 0.02 | <0.001 |
| GO:0016264 | gap junction assembly | 43 | 3 | 0.14 | <0.001 |
| GO:0045880 | positive regulation of smoothened signaling pathway | 107 | 4 | 0.34 | <0.001 |
| GO:0030828 | positive regulation of cGMP biosynthetic process | 50 | 3 | 0.16 | 0.001 |
| GO:0007406 | negative regulation of neuroblast proliferation | 51 | 3 | 0.16 | 0.001 |
| GO:0009103 | lipopolysaccharide biosynthetic process | 17 | 2 | 0.05 | 0.001 |
| GO:0007603 | phototransduction, visible light | 377 | 5 | 1.21 | 0.001 |
| GO:0007568 | aging | 1457 | 11 | 4.69 | 0.002 |
| GO:0060134 | prepulse inhibition | 84 | 3 | 0.27 | 0.003 |
| GO:0070474 | positive regulation of uterine smooth muscle contraction | 24 | 2 | 0.08 | 0.003 |
| GO:0043297 | apical junction assembly | 284 | 5 | 0.91 | 0.003 |
| GO:0051823 | regulation of synapse structural plasticity | 87 | 3 | 0.28 | 0.003 |
| GO:2001241 | positive regulation of extrinsic apoptotic in absence of ligand | 87 | 3 | 0.28 | 0.003 |
| GO:0010460 | positive regulation of heart rate | 139 | 4 | 0.45 | 0.004 |
| GO:0071504 | cellular response to heparin | 33 | 2 | 0.11 | 0.005 |
| GO:0035290 | trunk segmentation | 34 | 2 | 0.11 | 0.005 |
| GO:0043278 | response to morphine | 117 | 3 | 0.38 | 0.006 |
| GO:0006776 | vitamin A metabolic process | 38 | 2 | 0.12 | 0.007 |
| GO:0090136 | epithelial cell-cell adhesion | 122 | 3 | 0.39 | 0.007 |
| GO:0007210 | serotonin receptor signaling pathway | 40 | 2 | 0.13 | 0.007 |
| GO:0072047 | proximal/distal nephron patterning | 37 | 2 | 0.12 | 0.009 |
| *Polygenic selection* |  |  |  |  |  |
| GO:0030705 | cytoskeleton-dependent intracellular transport | 270 | 5 | 1.70 | <0.001 |
| GO:0051234 | establishment of localization | 6863 | 49 | 43.29 | 0.002 |
| GO:0043585 | nose morphogenesis | 14 | 2 | 0.09 | 0.003 |
| GO:0048870 | cell motility | 2926 | 23 | 18.45 | 0.005 |
| GO:0045163 | clustering of voltage-gated potassium channels | 17 | 2 | 0.11 | 0.005 |
| GO:0007195 | adenylate cyclase-inhibiting dopamine receptor signaling pathway | 17 | 2 | 0.11 | 0.005 |
| GO:0007166 | cell surface receptor signaling pathway | 4416 | 42 | 27.85 | 0.005 |
| GO:2001224 | positive regulation of neuron migration | 57 | 3 | 0.36 | 0.006 |
| GO:0010954 | positive regulation of protein processing | 18 | 2 | 0.11 | 0.006 |
| GO:0007616 | long-term memory | 120 | 4 | 0.76 | 0.007 |
| GO:0006979 | response to oxidative stress | 716 | 8 | 4.52 | 0.007 |
| GO:0007175 | negative regulation of epidermal growth factor-activated receptor activity | 21 | 2 | 0.13 | 0.008 |
| GO:0051489 | regulation of filopodium assembly | 182 | 5 | 1.15 | 0.009 |
| GO:0031547 | brain-derived neurotrophic factor receptor signaling pathway | 23 | 2 | 0.15 | 0.009 |
| GO:0051179 | localization | 8832 | 67 | 55.71 | 0.009 |

## Discussion

Overall, population genomic analysis of 728 juvenile Atlantic salmon collected throughout the Miramichi River system generated three primary conclusions. First, we have confirmed population structure of salmon in the Miramichi River is very weak compared with populations from other large river systems in North American and not hierarchical as would be expected for organisms inhabiting dendritic systems like rivers. Second, we provided one of the first insights into the genomic patterns related to a chromosomal fusion rearrangement. The fusion has previously been shown to be polymorphic in North American Atlantic salmon and we demonstrated that population structure for this chromosomal rearrangement is five times greater than the observed neutral population structure in the Miramichi River. The frequency of the fused chromosome further correlated with summer precipitation suggesting that variation in the presence of the fusion has functional importance for local adaptation of tributary populations. Third, we detected outlier variation among tributaries that was correlated with a gradient of elevation. Significantly, the detected outliers were clustered in a region on chromosome 9 that has been previously linked with important life history traits, including run-timing, in European populations of Atlantic salmon. These traits vary among tributaries in the Miramichi River and support the hypothesis that locally adaptive processes are more important than neutral processes for genetically structuring populations.

### Neutral population structure (or lack thereof?)

Large rivers containing populations of Atlantic salmon on both sides of the Atlantic Ocean frequently exhibit genetically distinct populations with levels of differentiation up to an order of magnitude greater than we report (e.g. Aykanat et al., 2015; Dionne et al., 2009; Primmer et al., 2006; Vähä et al., 2007). Strong philopatry and natal homing behaviours are defining traits of salmonids and are expected to result in hierarchical population structuring (Fraser et al., 2011). Our results revealed remarkably weak neutral population structure for the largest Atlantic salmon river in North America, the Miramichi River (Cunjak & Newbury, 2005). There was no evidence of hierarchical structuring of populations between the two main branches of the river and spatial structure only explained approximately 1% of the total allelic variation. Thus, the pattern of population structuring we have reported is at odds with expectations based on the species’ biology. While our results are consistent with two previous population genetic studies in the Miramichi River system based on a few microsatellites that also found weak structure and no support for hierarchical structuring of populations (Dionne et al., 2009; Moore et al., 2014), they contradict previous work based on transferrin gene allozyme variation (Møller, 1970). Allele frequency differences between Northwest and Southwest Miramichi River samples have been reported (Møller, 1970). The interpretation of Møller’s work is complicated because none of his samples from the Miramichi River conform to Hardy-Weinberg equilibrium expectations and spatial structure between Northwest and Southwest branches of the river is temporally confounded (spring vs. fall samples). The transferrin gene is located on chromosome 3 and although we detect outlier SNPs on this chromosome they occur tens of Mb from this gene’s location indicating a lack of association. As these data are derived from samples collected in the 1960s, it is possible that temporal changes in population structure have occurred that erased the historical population structure reported in his work.

Population divergence (F_ST_) is a function of the product of the effective population size and the number of migrants per generation (Allendorf & Phelps, 1981). Within the Miramichi River, two of the more divergent tributaries we identified had the lowest Nb estimates suggesting that higher genetic drift may explain their elevated neutral divergence in these cases. Yet, for the other tributaries, our adjusted Nb estimates were not larger than those reported in other Atlantic salmon rivers that exhibit stronger genetic divergence (Ferchaud et al., 2016; Palstra, O’Connell, & Ruzzante, 2007, 2009). These observations suggest that unusually large effective population sizes compared with other Atlantic salmon rivers are an unlikely explanation for the overall low levels of divergence we observed.

High rates of straying by spawning adults could explain the overall low divergence in the system. Straying for Atlantic salmon populations is typically at a rate of 6 – 10 % (Hendry, Castric, Kinnison, & Quinn, 2004; Keefer & Caudill, 2014; Pess, 2009); however, straying rates in the Miramichi River may be as high as 21% (Chaput, Moore, Hayward, Sheasgreen, & Dubee, 2001). What might cause naturally higher rates of straying in the Miramichi River compared to many other salmon rivers is unclear, but past stocking activities may play a role. The Miramichi River is stocked annually and has been for more than 100 years (Labadie, 2016) and salmonids reared in hatcheries can have straying rates higher than 20% (Keefer & Caudill, 2014). Current enhancement activities collect adults from specific tributaries and return offspring back to those tributaries (Labadie, 2016). These stocking efforts are believed to account for only a small fraction of contemporary adult returns (Chaput et al., 2001; Wallace & Curry, 2017) suggesting contemporary activities are an unlikely explanation; however, historical practises were not as restricted.

Stocking to enhance Atlantic salmon populations in the Miramichi River dates back to the 1870s. At times, as many as 4000 adult salmon broodstock are known to have been collected in the estuary, bred in the hatchery, and their offspring released throughout the watershed (M. Hambrook, Miramichi Salmon Association, pers. comm.). Depending on the frequency and duration of this stocking, these activities could give the appearance of highly elevated gene flow and may have erased historical signals of population structure. Stocking can disrupt the genetic structure of natural salmonid populations by reducing genetic divergence and increasing admixture among previously structured groups (Hansen, Fraser, Meier, & Mensberg, 2009; Marie, Bernatchez, & Garant, 2010; Pearse, Martinez, & Garza, 2011; Perrier, Baglinière, & Evanno, 2013; Perrier et al., 2011; Valiquette, Perrier, Thibault, & Bernatchez, 2014). Additionally, Atlantic salmon suffered a major anthropogenic insult in the 1950-60s, namely widespread pollution from DDT that resulted in mass mortalities (Elson, 1967; MacDonald, 1971), for which long-lasting effects on population structure are unclear. A combination of tagging studies, demographic modelling, and genotyping of archival samples to assess temporal trends in population structure would all be useful approaches to help resolve the underlying causes of weak population structuring in the Miramichi River.

For the three tributaries that show elevated divergence in this system, two of them exhibited small effective population sizes (Taxis River and Rocky Brook). In some years, Taxis River may be inaccessible to migrating adults due to low water levels (M. Hambrook, Miramichi Salmon Association, pers. comm.) suggesting that access to this system may drive the elevated genetic drift we observed. For the other two systems (Rocky Brook and Upper Northwest), there is no obvious reason why their connectivity to the rest of the system may be reduced. Both of these systems exhibited high alternate allele frequencies for the outlier region on chromosome 9 suggesting that selection may play a role in limiting the connectivity between these systems and the rest of the tributaries.

### Chromosomal rearrangement polymorphism: a possible role in local adaptation?

We provide the first molecular genetic characterization of the patterns of diversity for a polymorphic chromosomal rearrangement previously described in a North American Atlantic salmon aquaculture population (fusion between chromosome 8 and 29; Brenna-Hansen et al., 2012). Most notably, among-population variation in the frequency of the fusion did not mirror patterns of neutral variation, and rather co-varied with environmental variation in summer precipitation and elevation, suggesting a possible role in local adaptation. Different patterns of size-selective mortality on juveniles during summer have been demonstrated for Atlantic salmon under different precipitation regimes, where higher precipitation favoured smaller juveniles and lower precipitation favoured larger juveniles (Good, Dodson, Meekan, & Ryan, 2001). Several other studies in salmonids have also identified precipitation as an important variable underlying genomic variation among populations (Bourret, Dionne, et al., 2013; Hecht et al., 2015; Matala, Ackerman, Campbell, & Narum, 2014; Olsen et al., 2011). Further investigation is needed to properly test the link between the 8-29 fusion and phenotype/fitness in Atlantic salmon. Given the correlation with summer precipitation and elevation, we propose that the fusion may be associated with the ability to handle high flow velocity during the critical first few months of post-hatching juvenile salmon. While speculative, enriched gene functions detected in the fused region included cell-cell adhesion, smooth muscle contraction, and regulation of heart rate that could suggest circulatory system adaptations related to aerobic activity. Alternatively, this variation in precipitation could be related to triggering and the timing of migrations that differ among tributaries (Leggett, 1977). Other enriched gene functions detected in the fused region were related to neural development, phototransduction, aging, and behaviour that may indicate the fusion captured beneficial alleles related to variation in life history. It is further possible that the adaptive traits associated with the fusion are not captured in these enrichment tests due to the specifics of their genetic architecture.

Chromosomal rearrangements, in particular inversions, frequently display such polymorphism in natural populations, with variations of frequencies along environmental gradients, which is generally interpreted as correlative evidence for a role of the inversion in local adaptation (e.g. Kapun et al., 2016). In fact, inversions underlie important phenotypic and life history variation in a wide variety of organisms (Wellenreuther & Bernatchez, 2018). By limiting recombination between the standard and inverted rearrangements and by trapping together a set of alleles, inversions provide an important mechanism that facilitates local adaptation in the face of high levels of gene flow. In fish, inversions have been linked with a suite of traits differentiating anadromous and resident ecotypes of rainbow trout (Pearse, Miller, Abadia-Cardoso, & Garza, 2014), migratory phenotypes in Atlantic cod on both sides of the Atlantic Ocean (Berg et al., 2016; Sinclair-Waters et al., 2017), and adaptation to salinity gradients in Atlantic cod (Barth et al., 2017; Berg et al., 2015). Remarkably, the chromosomal rearrangement in cod allows fjord cod to maintain local adaptation to low salinity conditions despite high levels of immigration from offshore populations (Barth et al., 2017). Such mechanisms are also expected for fusions (Guerrero & Kirkpatrick, 2014) and may be particularly relevant for Atlantic salmon populations in the Miramichi River, which experience contrasting environments but weak neutral differentiation.

In contrast with the accumulation of studies on inversions, the role of fusions in adaptation is poorly documented. Karyotypic variation for fusions are widely reported within and between species (Dobigny et al., 2017) and for some taxa, such as the house mouse, can represent extreme levels of within-species karyotypic variation (Piálek, Hauffe, & Searle, 2005). Yet, very few studies report the distribution of fusion polymorphism in natural populations with possible links to fitness, phenotype, or the environment (Dobigny et al., 2017). Theoretical work suggests that chromosomal fusions may evolve in response to local selective forces (Guerrero & Kirkpatrick, 2014). To our knowledge, the only empirical support for this prediction is the grasshopper *Dichroplus pratensis*, for which fusion has been linked to phenotype and where fusion frequency varies with abiotic conditions, suggesting a link with adaptation to environmental extremes (Bidau, Miño, Castillo, & Martí, 2012). In such a context, the patterns of polymorphism and the environmental association we have observed for the chromosome 8 and 29 fusion in Miramichi River populations of Atlantic salmon represent a new opportunity to better understand whether, like inversions, fusions play a key role in adaptation in the face of gene flow. Recently work in review suggests the fusion between chromosomes 8 and 29 of Atlantic salmon is under balancing selection across its North American range providing independent evidence it has an adaptive role (Lehnert et al., 2018). The within-river structure we have demonstrated is consistent with this global pattern, and suggests a mechanism to explain the balancing selection detected by Lehnert et al. (2018). Frequency-dependent selection for different fusion-mediated life histories or spatially varying selection within rivers (e.g. upstream vs. downstream), such as we have demonstrated, would be expected to result in the genomic patterns of balancing selection at a wider scale. Importantly, variation maintained in this way by spatially balancing selection is believed to be more useful than other genetic mechanisms for maintaining variation that would allow populations to respond to future challenges (Whitlock, 2015).

The dense SNP dataset employed in our study also provides one of the first analyses of the genomic features associated with a chromosomal fusion and provides a complementary picture to previous knowledge based on cytology. For instance, patterns of allelic variation indicated that fusion homokaryotes exhibit much reduced diversity compared with the nonfusion homokaryotes. This result is consistent with meiotic observations that reported a reduced frequency of recombination events in fused chromosomes and a shift in the distribution of recombination toward the distal ends of chromosomes in fused homokaryotes and to a lesser extent in heterokaryotes (Bidau et al., 2001; Dumas & Britton-Davidian, 2002).

The patterns of linkage we report between these chromosomes also allow delimiting the regions affected by the reduction of recombination associated with the rearrangement. Previous work by Brenna-Hansen et al. (2012), which described this fusion, reported difficulties in determining the architecture of the fused chromosome from genetic mapping data. On the basis of fluorescence in-situ hybridization analysis, they had suggested that the fusion represented a tandem joining of the q-arm of chromosome 8 to the centromere of the acrocentric chromosome 29. Our linkage data suggests instead that it is the p-arm of chromosome 8 that has fused with the centromere of chromosome 29. The p-arm of chromosome 8 contains a dense cluster of rDNA repeats (Brenna-Hansen et al., 2012; Lien et al., 2016) and repeated elements of rDNA have been associated with chromosomal fusions and fissions in a wide variety of species (e.g. Barros et al., 2017; Cazaux, Catalan, Veyrunes, Douzery, & Britton-Davidian, 2011; Huang, Ma, Yang, Fei, & Li, 2008; Stahl et al., 1983) suggesting they play an important role in chromosomal rearrangements and thus may explain the occurrence of the fusion we have observed. Ultimately long-read sequencing data will be necessary to verify the architecture of this chromosomal rearrangement.

### Adaptive variation

In addition to the possible adaptive variation observed for the chromosome 8 and 29 fusion, we identified genetic distinctiveness along an environmental gradient of elevation for a likely candidate gene. Most of the outliers we detected were localized to a 250 Kb region of chromosome 9 that has previously been implicated in important life history variation for this species. In European Atlantic salmon, a particular a gene in this region, *six6*, has been associated with age-at-maturity in Atlantic salmon in Europe (Barson et al., 2015; Johnston et al., 2014), differences in run-timing within rivers, and local adaptation between tributaries (Cauwelier et al., 2017; Pritchard et al., 2018). Our study is the first to report direct evidence of within-river variation at this locus in a North American population of Atlantic salmon. Previously, Kusche et al. (2017) failed to find an association of six6-linked SNPs with age-at-maturity in four Atlantic salmon rivers from Québec. Their failure to repeat this association may have been a result of *six6* influencing a phenotype (e.g. size-at-maturity or run-timing) that is only correlated with age-at-maturity in certain rivers, as has been suggested by Barson et al. (2015).

In our study, variation in *six6* allele frequencies varied along an elevation gradient suggesting that run-timing, or a related trait, in the Miramichi River may too be influenced by variation at this locus. Run-timing in the Miramichi River has historically been bimodal with peaks that represent early (June – July) and late (Sept. – Nov.) returning fish respectively (Chaput, Douglas, & Hayward, 2016; Saunders, 1967). Adults spawning in the upper reaches of rivers return earlier and ascend higher than late returning salmon (Chaput et al., 2016). In particular, the fish from Rocky Brook are known to return early and they also exhibited the highest alternate allele frequencies for *six6* associated SNPs of any tributary. Fish in this river are morphologically distinct and exhibit earlier migration timing from those occupying lower reaches of the Miramichi River (Riddell & Leggett, 1981). They exhibit more fusiform bodies and longer paired fins consistent with life in higher velocity streams (Riddell & Leggett, 1981). These differences are heritable and persist when grown in common environments with salmon from lower elevation sites in the Miramichi River suggesting a genetic basis for these traits (Riddell et al., 1981). There is evidence of similar trait variation between salmon from the upper and lower sections of the NW Miramichi River (Elson, 1973). Despite the expected differences in body shape we did not detect differences in condition factor in the expected directions. Rocky Brook samples did not differ appreciably from downstream tributaries in this crude measure of shape and Upper Northwest samples were more robust than all other tributaries. This is in the opposite direction of predictions. Clearly more work is needed to link genomic variation with important phenotypic and life history variation in this system. In Europe, Pritchard et al. (Pritchard et al., 2018) found an association of the *six6* locus with a variable explaining “flow volume” for Atlantic salmon in the Teno/Tana River in Norway. Our results for a North American system thus suggest that stream parameters related to river size (flow volume and velocity) that differ between upper and lower reaches of large river systems may be mechanisms driving convergent local adaptations across continents and represents avenues for future study in relation to known differences in morphology and life history.

In addition to single locus tests, we employed a polygenic framework to investigate correlated allelic changes among tributaries to environmental predictors. The same temperature / elevation variable was associated with correlated changes in allele frequency for 198 SNPs including 5 of those previously identified on chromosome 9. Gene function enrichment revealed roles in morphological development, memory, and sexual maturation for these genes that are consistent with the previously discussed differences in life history between tributaries. In total, nine of our variants detected by the RDA were located in proximity (<10 Kb) to variants associated with flow volume by Pritchard et al. (2018). This represented 14% of the flow-volume-associated SNPs detected by Pritchard et al. (2018) suggesting there is a level of genomic parallelism across continents. Of note, we too identified the neuronal growth regulator 1 gene (negr-1) in our temperature / elevation associated genes that has previously been suggested as a candidate locus for further investigation of its role in salmonid life history variation (Pritchard et al., 2018).

### Management implications

Our data have soundly rejected the presence of hierarchical structuring of Atlantic salmon populations in the Miramichi River that had been previously hypothesized. Instead our results support the relative isolation of upper watersheds from those in the lower river and a primary role for adaptive processes in structuring populations in this system. Furthermore, our work illustrates the importance of moving beyond single locus variation for investigating the occurrence of local adaptation at the genome level. Structural variation, such as the fusion variation we have characterized, is increasingly recognized for its role in maintaining life history and adaptive variation (Wellenreuther & Bernatchez, 2018). Thus, basing management decisions solely on results of single locus analyses may over-look important signatures of variation and generate sub-optimal recommendations for management. Our results have important relevance for the management of Atlantic salmon in the Miramichi River. Given the recent trend of run-timing homogenization (Douglas, Chaput, Hayward, & Sheasgreen, 2015), more attention must be paid to the underlying mechanisms generating life history variation and their role in population declines in this system. Ongoing and proposed supplementation activities to enhance populations must also continue to respect the potential local adaptation of tributary populations.

### Conclusions

We have generated the largest and highest resolution dataset for Atlantic salmon in the Miramichi River to date. We reported unexpectedly weak neutral population structure and a complete absence of support for hierarchical structuring in this large river system. In contrast, we characterized patterns of genomic variation for a polymorphic chromosomal fusion and demonstrated this genomic region exhibits significant differentiation among tributary populations. Our analyses further revealed associations between fusion frequency and environmental parameters orthogonal to that of other genomic variation, suggesting that this fusion rearrangement may contribute to local adaptation in the face of gene flow. We also characterized outlier SNPs whose frequencies correlated with environmental attributes and was consistent with both known life-history variation in this and in multiple other systems throughout the species’ range. Our results contribute to a growing body of literature implicating parallel selection at the *six6* gene for explaining within-river local adaptation of Atlantic salmon on both sides of the Atlantic Ocean. Our results represent significant progress toward the resolution of a long-standing debate over the presence of genetic structure for Atlantic salmon in the Miramichi River. They reveal that adaptive processes appear to be more important for structuring populations than neutral processes. Whether this reflects intrinsic properties of the populations (i.e. large effective population size) or a legacy of anthropogenic influence is an open and important question that is in urgent need of resolution for the improved conservation and management of this species.

## Supporting information

## Acknowledgements

This project would not have been possible without support from the Collaboration for Atlantic Salmon Tomorrow (CAST) and its partners, Cooke Aquaculture Inc. and JD Irving Ltd. This project was supported partially by a financial contribution from Fisheries and Oceans Canada. Sampling was coordinated by the Miramichi Salmon Association (Mark Hambrook and Holly Labadie) and Fisheries and Oceans Canada (Gerald Chaput and Scott Douglas). We also acknowledge staff at the Centre for Aquaculture Technologies Canada, PEI for DNA extraction services and staff at CIGENE, Ås, Norway for conducting the SNP genotyping. We thank Maren Wellenreuther, Julia Barth, and an anonymous reviewer for their constructive comments that improved an earlier version of this manuscript.

## Data Accessibility

The SNP genotype data in PLINK format and the environmental data for sites will be available on Dryad following article acceptance. Scripts used to analyze the data are available as a GitHub repository: https://github.com/kylewellband/ssa_cast2016.

## Author Contributions

TL, AC, and LB conceived the study, TL organized sample collection, JE provided the SNP genotyping platform, KW and CM analyzed the data, KW wrote the first draft, and all authors contributed to the final version.

## Literature Cited

Alexa, A., Rahnenführer, J., & Lengauer, T. (2006). Improved scoring of functional groups from gene expression data by decorrelating GO graph structure. Bioinformatics, 22(13), 1600–1607. doi:10.1093/bioinformatics/btl140

Alexander, D. H., Novembre, J., & Lange, K. (2009). Fast model-based estimation of ancestry in unrelated individuals. Genome Research, 19(9), 1655–1664. doi:10.1101/gr.094052.109

Allendorf, F. W., & Phelps, S. R. (1981). Use of Allelic Frequencies to Describe Population Structure. Canadian Journal of Fisheries and Aquatic Sciences, 38(12), 1507–1514. doi:10.1139/f81-203

Aykanat, T., Johnston, S. E., Orell, P., Niemelä, E., Erkinaro, J., & Primmer, C. R. (2015). Low but significant genetic differentiation underlies biologically meaningful phenotypic divergence in a large Atlantic salmon population. Molecular Ecology, 24(20), 5158–5174. doi:10.1111/mec.13383

Aykanat, T., Lindqvist, M., Pritchard, V. L., & Primmer, C. R. (2016). From population genomics to conservation and management: a workflow for targeted analysis of markers identified using genome-wide approaches in Atlantic salmon Salmo salar. Journal of Fish Biology, 89(6), 2658–2679. doi:10.1111/jfb.13149

Barros, A. V., Wolski, M. A. V., Nogaroto, V., Almeida, M. C., Moreira-Filho, O., & Vicari, M. R. (2017). Fragile sites, dysfunctional telomere and chromosome fusions: What is 5S rDNA role? Gene, 608, 20–27. doi:10.1016/j.gene.2017.01.013

Barson, N. J., Aykanat, T., Hindar, K., Baranski, M., Bolstad, G. H., Fiske, P., … Primmer, C. R. (2015). Sex-dependent dominance at a single locus maintains variation in age at maturity in salmon. Nature, 528(7582), 405–408. doi:10.1038/nature16062

Barth, J. M. I., Berg, P. R., Jonsson, P. R., Bonanomi, S., Corell, H., Hemmer-Hansen, J., … André, C. (2017). Genome architecture enables local adaptation of Atlantic cod despite high connectivity. Molecular Ecology, 26(17), 4452–4466. doi:10.1111/mec.14207

Benestan, L., Quinn, B. K., Maaroufi, H., Laporte, M., Clark, F. K., Greenwood, S. J., … Bernatchez, L. (2016). Seascape genomics provides evidence for thermal adaptation and current-mediated population structure in American lobster (Homarus americanus). Molecular Ecology, 25(20), 5073–5092. doi:10.1111/mec.13811

Benjamini, Y., & Hochberg, Y. (1995). Controlling the False Discovery Rate: A Practical and Powerful Approach to Multiple Testing. Journal of the Royal Statistical Society B, 57(1), 289–300.

Berg, P. R., Jentoft, S., Star, B., Ring, K. H., Knutsen, H., Lien, S., … André, C. (2015). Adaptation to Low Salinity Promotes Genomic Divergence in Atlantic Cod (Gadus morhua L.). Genome Biology and Evolution, 7(6), 1644–1663. doi:10.1093/gbe/evv093

Berg, P. R., Star, B., Pampoulie, C., Sodeland, M., Barth, J. M. I., Knutsen, H., … Jentoft, S. (2016). Three chromosomal rearrangements promote genomic divergence between migratory and stationary ecotypes of Atlantic cod. Scientific Reports, 6(1), 23246. doi:10.1038/srep23246

Bidau, C. J., Giménez, M. D., Palmer, C. L., & Searle, J. B. (2001). The effects of Robertsonian fusions on chiasma frequency and distribution in the house mouse (Mus musculus domesticus) from a hybrid zone in northern Scotland. Heredity, 87(3), 305–313. doi:10.1046/j.1365-2540.2001.00877.x

Bidau, C. J., Miño, C. I., Castillo, E. R., & Martí, D. A. (2012). Effects of Abiotic Factors on the Geographic Distribution of Body Size Variation and Chromosomal Polymorphisms in Two Neotropical Grasshopper Species (Dichroplus : Melanoplinae: Acrididae). Psyche: A Journal of Entomology, 2012, 1–11. doi:10.1155/2012/863947

Bonhomme, M., Chevalet, C., Servin, B., Boitard, S., Abdallah, J., Blott, S., & SanCristobal, M. (2010). Detecting selection in population trees: The Lewontin and Krakauer test extended. Genetics, 186(1), 241–262. doi:10.1534/genetics.110.117275

Bourret, V., Dionne, M., Kent, M. P., Lien, S., & Bernatchez, L. (2013). Landscape genomics in Atlantic salmon (salmo salar): Searching for gene-environment interactions driving local adaptation. Evolution, 67(12), 3469–3487. doi:10.1111/evo.12139

Bourret, V., Kent, M. P., Primmer, C. R., Vasemägi, A., Karlsson, S., Hindar, K., … Lien, S. (2013). SNP-array reveals genome-wide patterns of geographical and potential adaptive divergence across the natural range of Atlantic salmon (Salmo salar). Molecular Ecology, 22(3), 532–551. doi:10.1111/mec.12003

Brauer, C. J., Unmack, P. J., Smith, S., Bernatchez, L., & Beheregaray, L. B. (2018). On the roles of landscape heterogeneity and environmental variation in determining population genomic structure in a dendritic system. Molecular Ecology, (March), 3484–3497. doi:10.1111/mec.14808

Brenna-Hansen, S., Li, J., Kent, M. P., Boulding, E. G., Dominik, S., Davidson, W. S., & Lien, S. (2012). Chromosomal differences between European and North American Atlantic salmon discovered by linkage mapping and supported by fluorescence in situ hybridization analysis. BMC Genomics, 13(1). doi:10.1186/1471-2164-13-432

Cáceres, A., & González, J. R. (2015). Following the footprints of polymorphic inversions on SNP data: from detection to association tests. Nucleic Acids Research, 43(8), e53–e53. doi:10.1093/nar/gkv073

Cáceres, A., Sindi, S. S., Raphael, B. J., Cáceres, M., & González, J. R. (2012). Identification of polymorphic inversions from genotypes. BMC Bioinformatics, 13, 28. doi:10.1186/1471-2105-13-28

Cauwelier, E., Gilbey, J., Sampayo, J., Stradmeyer, L., & Middlemas, S. J. (2017). Identification of a single genomic region associated with seasonal river return timing in adult Scottish Atlantic salmon (Salmo salar L.) identified using a genome-wide association study. Canadian Journal of Fisheries and Aquatic Sciences, cjfas-2017-0293. doi:10.1139/cjfas-2017-0293

Cazaux, B., Catalan, J., Veyrunes, F., Douzery, E. J., & Britton-Davidian, J. (2011). Are ribosomal DNA clusters rearrangement hotspots? A case study in the genus Mus (Rodentia, Muridae). BMC Evolutionary Biology, 11(1), 124. doi:10.1186/1471-2148-11-124

Chang, C. C., Chow, C. C., Tellier, L. C. A. M., Vattikuti, S., Purcell, S. M., & Lee, J. J. (2015). Second-generation PLINK: Rising to the challenge of larger and richer datasets. GigaScience, 4(1), 1–16. doi:10.1186/s13742-015-0047-8

Chaput, G., Douglas, S. G., & Hayward, J. (2016). Biological Characteristics and Population Dynamics of Atlantic Salmon (Salmo salar) from the Miramichi River, New Brunswick, Canada. Canadian Science Advisory Secretariat Research Document 2016/029.

Chaput, G., Moore, D., Hayward, J., Sheasgreen, J., & Dubee, B. (2001). Stock status of Atlantic Salmon (Salmo salar) in the Miramichi River, 2000. Canadian Science Advisory Secretariat Research Document 2001/008, 1–88.

Cunjak, R. A., & Newbury, R. W. (2005). Atlantic Coast Rivers of Canada. In A. C. Benke & C. E. Cushing (Eds.), Rivers of North America (pp. 939–980). San Diego, CA: Elsevier Inc. (Academic Press).

DFO. (2017). Update of indicators to 2017 of adult Altantic salmon for the Miramichi River (NB), Salmon Fishing Area 16, DFO Gulf Region. Canadian Science Advisory Secretariat Science Response 2017/043, 1–7.

Dionne, M., Caron, F., Dodson, J. J., & Bernatchez, L. (2008). Landscape genetics and hierarchical genetic structure in Atlantic salmon: The interaction of gene flow and local adaptation. Molecular Ecology, 17(10), 2382–2396. doi:10.1111/j.1365-294X.2008.03771.x

Dionne, M., Caron, F., Dodson, J. J., & Bernatchez, L. (2009). Comparative survey of within-river genetic structure in Atlantic salmon; relevance for management and conservation. Conservation Genetics, 10(4), 869–879. doi:10.1007/s10592-008-9647-5

Dionne, M., Miller, K. M., Dodson, J. J., Caron, F., & Bernatchez, L. (2007). Clinal variation in MHC diversity with temperature: Evidence for the role of host-pathogen interaction on local adaptation in Atlantic salmon. Evolution, 61(9), 2154–2164. doi:10.1111/j.1558-5646.2007.00178.x

Do, C., Waples, R. S., Peel, D., Macbeth, G. M., Tillett, B. J., & Ovenden, J. R. (2014). NeEstimator v2: Re-implementation of software for the estimation of contemporary effective population size (Ne) from genetic data. Molecular Ecology Resources, 14(1), 209–214. doi:10.1111/1755-0998.12157

Dobigny, G., Britton-Davidian, J., & Robinson, T. J. (2017). Chromosomal polymorphism in mammals: an evolutionary perspective. Biological Reviews, 92(1), 1–21. doi:10.1111/brv.12213

Dodson, J., & Colombani, F. (1997). The genetic identity of the Clearwater Brook population of Atlantic salmon (Salmo salar); a temporal and spatial study of Atlantic salmon population genetic structure in the Miramichi, St. John and Margaree Rivers. Atlantic Salmon Federation Report, 1–24.

Douglas, S. G., Chaput, G., Hayward, J., & Sheasgreen, J. (2015). Assessment of Atlantic Salmon (Salmo salar) in Salmon Fishing Area 16 of the southern Gulf of St. Lawrence to 2013. Canadian Science Advisory Secretariat Research Document 2015/049. Retrieved from http://www.dfo-mpo.gc.ca/csas-sccs/

Dray, S., Blanchet, G., Borcard, D., Clappe, S., Guenard, G., Jombart, T., Larocque, G., Legendre, P., Madi, N., & Wagner, H. (2018). adespatial: Multivariate Multiscale Spatial Analysis. R package version 0.1-1. https://CRAN.R-project.org/package=adespatial

Dumas, D., & Britton-Davidian, J. (2002). Chromosomal rearrangements and evolution of recombination: Comparison of chiasma distribution patterns in standard and Robertsonian populations of the house mouse. Genetics, 162(3), 1355–1366.

Elson, P. F. (1967). Effects on Wild Young Salmon of Spraying DDT over New Brunswick Forests. Journal of the Fisheries Research Board of Canada, 24(4), 731–767. doi:10.1139/f67-066

Elson, P. F. (1973). Genetic polymorphism in Northwest Miramishi salmon, in relation to season of river ascent and age at maturation and its implications for management of the stocks. International Commission for Northwest Atlantic Fisheries Research Document, 73(76).

Excoffier, L., & Lischer, H. E. L. (2010). Arlequin suite ver 3.5: A new series of programs to perform population genetics analyses under Linux and Windows. Molecular Ecology Resources, 10(3), 564–567. doi:10.1111/j.1755-0998.2010.02847.x

Ferchaud, A. L., Perrier, C., April, J., Hernandez, C., Dionne, M., & Bernatchez, L. (2016). Making sense of the relationships between Ne, Nb and Nc towards defining conservation thresholds in Atlantic salmon (Salmo salar). Heredity, 117(4), 268–278. doi:10.1038/hdy.2016.62

Fick, S. E., & Hijmans, R. J. (2017). WorldClim 2: new 1-km spatial resolution climate surfaces for global land areas. International Journal of Climatology, 37(12), 4302–4315. doi:10.1002/joc.5086

Foll, M., & Gaggiotti, O. (2008). A genome-scan method to identify selected loci appropriate for both dominant and codominant markers: A Bayesian perspective. Genetics, 180(2), 977–993. doi:10.1534/genetics.108.092221

Forester, B. R., Lasky, J. R., Wagner, H. H., & Urban, D. L. (2018). Comparing methods for detecting multilocus adaptation with multivariate genotype-environment associations. Molecular Ecology, 27(9), 2215–2233. doi:10.1111/mec.14584

Fourcade, Y., Chaput-Bardy, A., Secondi, J., Fleurant, C., & Lemaire, C. (2013). Is local selection so widespread in river organisms? Fractal geometry of river networks leads to high bias in outlier detection. Molecular Ecology, 22(8), 2065–2073. doi:10.1111/mec.12158

Fraser, D. J., & Bernatchez, L. (2001). Adaptive evolutionary conservation: towards a unified concept for defining conservation units. Molecular Ecology, 10(12), 2741–52. Retrieved from http://www.ncbi.nlm.nih.gov/pubmed/11903888

Fraser, D. J., Weir, L. K., Bernatchez, L., Hansen, M. M., & Taylor, E. B. (2011). Extent and scale of local adaptation in salmonid fishes: review and meta-analysis. Heredity, 106(3), 404–420. doi:10.1038/hdy.2010.167

Frontier, S. (1976). Étude de la décroissance des valeurs propres dans une analyse en composantes principales: Comparaison avec le moddle du bâton brisé. Journal of Experimental Marine Biology and Ecology, 25(1), 67–75. doi:10.1016/0022-0981(76)90076-9

Funk, W. C., McKay, J. K., Hohenlohe, P. A., & Allendorf, F. W. (2012). Harnessing genomics for delineating conservation units. Trends in Ecology and Evolution, 27(9), 489–496. doi:10.1016/j.tree.2012.05.012

Garant, D., Dodson, J. J., & Bernatchez, L. (2000). Ecological determinants and temporal stability of the within-river population structure in Atlantic salmon (Salmo salar L.). Molecular Ecology, 9(5), 615–628. doi:10.1046/j.1365-294X.2000.00909.x

Garcia de Leaniz, C., Fleming, I. A., Einum, S., Verspoor, E., Jordan, W. C., Consuegra, S., … Quinn, T. P. (2007). A critical review of adaptive genetic variation in Atlantic salmon: implications for conservation. Biological Reviews, 82(2), 173–211. doi:10.1111/j.1469-185X.2006.00004.x

Gautier, M. (2015). Genome-Wide Scan for Adaptive Divergence and Association with Population-Specific Covariates. Genetics, 201(4), 1555–1579. doi:10.1534/genetics.115.181453

Good, S. P., Dodson, J. J., Meekan, M. G., & Ryan, D. A. J. (2001). Annual variation in size-selective mortality of Atlantic salmon (*Salmo salar*) fry. Canadian Journal of Fisheries and Aquatic Sciences, 58(6), 1187–1195. doi:10.1139/cjfas-58-6-1187

Guerrero, R. F., & Kirkpatrick, M. (2014). Local adaptation and the evolution of chromosome fusions. Evolution, 68(10), 2747–2756. doi:10.1111/evo.12481

Hansen, M. M., Fraser, D. J., Meier, K., & Mensberg, K.-L. D. (2009). Sixty years of anthropogenic pressure: a spatio-temporal genetic analysis of brown trout populations subject to stocking and population declines. Molecular Ecology, 18(12), 2549–2562. doi:10.1111/j.1365-294X.2009.04198.x

Hartigan, J. A., & Hartigan, P. M. (1985). The dip test for unimodalty. The Annals of Statistics, 13(1), 70–84.

Hecht, B. C., Matala, A. P., Hess, J. E., & Narum, S. R. (2015). Environmental adaptation in Chinook salmon (Oncorhynchus tshawytscha) throughout their North American range. Molecular Ecology, 24(22), 5573–5595. doi:10.1111/mec.13409

Hendry, A. P., Castric, V., Kinnison, M. T., & Quinn, T. P. (2004). The Evolution of Philopatry and Dispersal: Homing Versus Straying in Salmonids. In A. P. Hendry & S. C. Stearns (Eds.), Evolution Illuminated: Salmon and thier relatives (pp. 52–91). New York, NY: Oxford University Press.

Huang, J., Ma, L., Yang, F., Fei, S. Z., & Li, L. (2008). 45S rDNA regions are chromosome fragile sites expressed as gaps in vitro on metaphase chromosomes of root-tip meristematic cells in Lolium spp. PLoS ONE, 3(5), 1–7. doi:10.1371/journal.pone.0002167

Jeffery, N. W., Stanley, R. R. E., Wringe, B. F., Guijarro-Sabaniel, J., Bourret, V., Bernatchez, L., … Bradbury, I. R. (2017). Range-wide parallel climate-associated genomic clines in Atlantic salmon. Royal Society Open Science, 4(11), 171394. doi:10.1098/rsos.171394

Johnston, S. E., Orell, P., Pritchard, V. L., Kent, M. P., Lien, S., Niemelä, E., … Primmer, C. R. (2014). Genome-wide SNP analysis reveals a genetic basis for sea-age variation in a wild population of Atlantic salmon (Salmo salar). Molecular Ecology, 23(14), 3452–3468. doi:10.1111/mec.12832

Jombart, T., & Ahmed, I. (2011). adegenet 1.3-1: New tools for the analysis of genome-wide SNP data. Bioinformatics, 27(21), 3070–3071. doi:10.1093/bioinformatics/btr521

Jombart, T., Devillard, S., & Balloux, F. (2010). Discriminant analysis of principal components: a new method for the analysis of genetically structured populations. BMC Genetics, 11(1), 94. doi:10.1186/1471-2156-11-94

Kapun, M., Fabian, D. K., Goudet, J., & Flatt, T. (2016). Genomic Evidence for Adaptive Inversion Clines in Drosophila melanogaster. Molecular Biology and Evolution, 33(5), 1317–1336. doi:10.1093/molbev/msw016

Keefer, M. L., & Caudill, C. C. (2014). Homing and straying by anadromous salmonids: A review of mechanisms and rates. Reviews in Fish Biology and Fisheries, 24(1), 333–368. doi:10.1007/s11160-013-9334-6

Kusche, H., Côté, G., Hernandez, C., Normandeau, E., Boivin-Delisle, D., & Bernatchez, L. (2017). Characterization of natural variation in North American Atlantic Salmon populations (Salmonidae: Salmo salar) at a locus with a major effect on sea age. Ecology and Evolution, 7(15), 5797–5807. doi:10.1002/ece3.3132

Labadie, H. (2016). Miramichi Salmon and Trout Restoration - Stocking 2016.

Leggett, W. C. (1977). The ecology of fish migrations. Annual Review of Ecology and Systematics, (8), 285–308.

Lehnert, S. J., Bentzen, P., Kess, T., Lien, S., Horne, J. B., Clement, M., & Bradbury, I. R. (2018). Chromosome polymorphisms track trans-Atlantic divergence, admixture and adaptive evolution in salmon. BioRxiv, 351338. doi:10.1101/351338

Lien, S., Koop, B. F., Sandve, S. R., Miller, J. R., Kent, M. P., Nome, T., … Davidson, W. S. (2016). The Atlantic salmon genome provides insights into rediploidization. Nature, 533(7602), 200–205. doi:10.1038/nature17164

Lindtke, D., Lucek, K., Soria-Carrasco, V., Villoutreix, R., Farkas, T. E., Riesch, R., … Nosil, P. (2017). Long-term balancing selection on chromosomal variants associated with crypsis in a stick insect. Molecular Ecology, 26(22), 6189–6205. doi:10.1111/mec.14280

Lotterhos, K. E., & Whitlock, M. C. (2014). Evaluation of demographic history and neutral parameterization on the performance of FST outlier tests. Molecular Ecology, 23(9), 2178–2192. doi:10.1111/mec.12725

Luu, K., Bazin, E., & Blum, M. G. B. (2017). pcadapt : an R package to perform genome scans for selection based on principal component analysis. Molecular Ecology Resources, 17(1), 67–77. doi:10.1111/1755-0998.12592

MacDonald, J. R. (1971). Delayed mortality of Atlantic salmon (Salmo salar) parr after forest spraying with DDT. Canadian Fisheries and Forestry Resource Development Branch Progress Report No. 2, 1–26.

Marie, A. D., Bernatchez, L., & Garant, D. (2010). Loss of genetic integrity correlates with stocking intensity in brook charr (Salvelinus fontinalis). Molecular Ecology, 19(10), 2025–2037. doi:10.1111/j.1365-294X.2010.04628.x

Matala, A. P., Ackerman, M. W., Campbell, M. R., & Narum, S. R. (2014). Relative contributions of neutral and non-neutral genetic differentiation to inform conservation of steelhead trout across highly variable landscapes. Evolutionary Applications, 7(6), 682–701. doi:10.1111/eva.12174

Meirmans, P. G. (2015). Seven common mistakes in population genetics and how to avoid them. Molecular Ecology, 24(13), 3223–3231. doi:10.1111/mec.13243

Mobley, K. B., Granroth-wilding, H., Ellmen, M., Vähä, J., Johnston, S. E., Orell, P., … Primmer, C. R. (2018). Home ground advantage : selection against dispersers promotes cryptic local adaptation in wild salmon. BioRxiv. doi:10.1101/311258

Møller, D. (1970). Transferrin Polymorphism in Atlantic Salmon (Salmo salar). Journal of the Fisheries Research Board of Canada, 27(9), 1617–1625. doi:10.1139/f70-182

Møller, D. (2005). Genetic studies on serum transferrins in Atlantic salmon. Journal of Fish Biology, 67(s1), 55–67. doi:10.1111/j.0022-1112.2005.00839.x

Moore, J. S., Bourret, V., Dionne, M., Bradbury, I., O’Reilly, P., Kent, M., … Bernatchez, L. (2014). Conservation genomics of anadromous Atlantic salmon across its North American range: Outlier loci identify the same patterns of population structure as neutral loci. Molecular Ecology, 23(23), 5680–5697. doi:10.1111/mec.12972

NRCAN (2016). Canada Digital Elevation Model, Edition 1.1. https://open.canada.ca/data/dataset/7f245e4d-76c2-4caa-951a-45d1d2051333. Accessed: February 16, 2018.

Oksanen, J., Blanchet, F. G., Friendly, M., Kindt, R., Legendre, P., McGlinn, D., Minchin, P. R., O’Hara, R. B., Simpson, G. L., Solymos, P., Stevens, M. H. H., Szoecs E., & Wagner H. (2017). vegan: Community Ecology Package. R package version 2.4-5. https://CRAN.R-project.org/package=vegan

Olsen, J. B., Crane, P. A., Flannery, B. G., Dunmall, K., Templin, W. D., & Wenburg, J. K. (2011). Comparative landscape genetic analysis of three Pacific salmon species from subarctic North America. Conservation Genetics, 12(1), 223–241. doi:10.1007/s10592-010-0135-3

Palsbøll, P. J., Bérubé, M., & Allendorf, F. W. (2007). Identification of management units using population genetic data. Trends in Ecology and Evolution, 22(1), 11–16. doi:10.1016/j.tree.2006.09.003

Palstra, F. P., O’Connell, M. F., & Ruzzante, D. E. (2007). Population structure and gene flow reversals in Atlantic salmon (Salmo salar) over contemporary and long-term temporal scales: Effects of population size and life history. Molecular Ecology, 16(21), 4504–4522. doi:10.1111/j.1365-294X.2007.03541.x

Palstra, F. P., O’Connell, M. F., & Ruzzante, D. E. (2009). Age structure, changing demography and effective population size in Atlantic salmon (Salmo salar). Genetics, 182(4), 1233–1249. doi:10.1534/genetics.109.101972

Patterson, N., Price, A. L., & Reich, D. (2006). Population Structure and Eigenanalysis. PLoS Genetics, 2(12), e190. doi:10.1371/journal.pgen.0020190

Pearse, D. E., Martinez, E., & Garza, J. C. (2011). Disruption of historical patterns of isolation by distance in coastal steelhead. Conservation Genetics, 12(3), 691–700. doi:10.1007/s10592-010-0175-8

Pearse, D. E., Miller, M. R., Abadia-Cardoso, A., & Garza, J. C. (2014). Rapid parallel evolution of standing variation in a single, complex, genomic region is associated with life history in steelhead/rainbow trout. Proceedings of the Royal Society B: Biological Sciences, 281(1783), 20140012–20140012. doi:10.1098/rspb.2014.0012

Pendás, A. M., Morán, P., & Garcia-Vázquez, E. (1993). Ribosomal RNA genes are interspersed throughout a heterochromatic chromosome arm in Atlantic salmon. Cytogenetic and Genome Research, 63(2), 128–130. doi:10.1159/10.1159/000133517

Perrier, C., Baglinière, J. L., & Evanno, G. (2013). Understanding admixture patterns in supplemented populations: A case study combining molecular analyses and temporally explicit simulations in Atlantic salmon. Evolutionary Applications, 6(2), 218–230. doi:10.1111/j.1752-4571.2012.00280.x

Perrier, C., Guyomard, R., Bagliniere, J.-L., & Evanno, G. (2011). Determinants of hierarchical genetic structure in Atlantic salmon populations: environmental factors vs. anthropogenic influences. Molecular Ecology, 20(20), 4231–4245. doi:10.1111/j.1365-294X.2011.05266.x

Pess, G. R. (2009). Patterns and Processes of Salmon Colonization. University of Washington, Seattle WA.

Piálek, J., Hauffe, H. C., & Searle, J. B. (2005). Chromosomal variation in the house mouse. Biological Journal of the Linnean Society, 84(3), 535–563. doi:10.1111/j.1095-8312.2005.00454.x

Primmer, C. R., Veselov, A. J., Zubchenko, A., Poututkin, A., Bakhmet, I., & Koskinen, M. T. (2006). Isolation by distance within a river system: Genetic population structuring of Atlantic salmon, Salmo salar, in tributaries of the Varzuga River in northwest Russia. Molecular Ecology, 15(3), 653–666. doi:10.1111/j.1365-294X.2005.02844.x

Pritchard, V. L., Mäkinen, H., Vähä, J.-P., Erkinaro, J., Orell, P., & Primmer, C. R. (2018). Genomic signatures of fine-scale local selection in Atlantic salmon suggest involvement of sexual maturation, energy homeostasis and immune defence-related genes. Molecular Ecology, 27(11), 2560–2575. doi:10.1111/mec.14705

QGIS Development Team (2016). QGIS Geographic Information System. Open Source Geospatial Foundation Project. http://qgis.osgeo.org

Quinlan, A. R., & Hall, I. M. (2010). BEDTools: a flexible suite of utilities for comparing genomic features. Bioinformatics, 26(6), 841–842. doi:10.1093/bioinformatics/btq033

R Core Team (2017). R: A language and environment for statistical computing. R Foundation for Statistical Computing, Vienna, Austria. https://www.R-project.org

Riddell, B. E., & Leggett, W. C. (1981). Evidence of an Adaptive Basis for Geographic Variation in Body Morphology and Time of Downstream Migration of Juvenile Atlantic Salmon (*Salmo salar*). Canadian Journal of Fisheries and Aquatic Sciences, 38(3), 308–320. doi:10.1139/f81-042

Riddell, B. E., Leggett, W. C., & Saunders, R. L. (1981). Evidence of adaptive polygenic variation between two populations of Atlantic salmon (Salmo salar) native to tributaries of the SW Miramichi River, N. B. Canandian Journal of Fisheries and Aquatic Science, 38(5605 1), 321–333.

Samy, J. K. A., Mulugeta, T. D., Nome, T., Sandve, S. R., Grammes, F., Kent, M. P., … Våge, D. I. (2017). SalmoBase: An integrated molecular data resource for Salmonid species. BMC Genomics, 18(1), 1–5. doi:10.1186/s12864-017-3877-1

Saunders, R. L. (1967). Seasonal Pattern of Return of Atlantic Salmon in the Northwest Miramichi River, New Brunswick. Journal of the Fisheries Research Board of Canada, 24(1), 21–32. doi:10.1139/f67-003

Schaffer, W. M., & Elson, P. F. (1975). The Adaptive Significance of Variations in Life History among Local Populations of Atlantic Salmon in North America. Ecology, 56(3), 577–590. Retrieved from https://www.jstor.org/stable/1935492

Sinclair-Waters, M., Bradbury, I. R., Morris, C. J., Lien, S., Kent, M. P., & Bentzen, P. (2017). Ancient chromosomal rearrangement associated with local adaptation of a postglacially colonized population of Atlantic Cod in the northwest Atlantic. Molecular Ecology, (November), 1–13. doi:10.1111/mec.14442

Stahl, A., Luciani, J. M., Hartung, M., Devictor, M., Bergé-Lefranc, J. L., & Guichaoua, M. (1983). Structural basis for Robertsonian translocations in man: association of ribosomal genes in the nucleolar fibrillar center in meiotic spermatocytes and oocytes. Proceedings of the National Academy of Sciences of the United States of America, 80(19), 5946–5950. doi:10.1073/pnas.80.19.5946

Swansburg, E., Chaput, G., Moore, D., Caissie, D., & El-Jabi, N. (2002). Size variability of juvenile Atlantic salmon: Links to environmental conditions. Journal of Fish Biology, 61(3), 661–683. doi:10.1006/jfbi.2002.2088

Tyers, M. (2017). riverdist: River Network Distance Computation and Applications. R package version 0.15.0. https://CRAN.R-project.org/package=riverdist

Vähä, J.-P., Erkinaro, J., Niemelä, E., & Primmer, C. R. (2007). Life-history and habitat features influence the within-river genetic structure of Atlantic salmon. Molecular Ecology, 16(13), 2638–2654. doi:10.1111/j.1365-294X.2007.03329.x

Valiquette, E., Perrier, C., Thibault, I., & Bernatchez, L. (2014). Loss of genetic integrity in wild lake trout populations following stocking: insights from an exhaustive study of 72 lakes from Québec, Canada. Evolutionary Applications, 7(6), 625–644. doi:10.1111/eva.12160

Verspoor, E., Beardmore, J. A., Consuegra, S., Garcia de Leaniz, C., Hindar, K., Jordan, W. C., … Cross, T. F. (2005). Population structure in the Atlantic salmon: insights from 40 years of research into genetic protein variation. Journal of Fish Biology, 67, 3–54. doi:10.1111/j.1095-8649.2005.00838.x

Wallace, B. G., & Curry, R. A. (2017). Assessing the outcomes of stocking hatchery-reared juveniles in the presence of wild Atlantic salmon. Environmental Biology of Fishes, 100(7), 877–887. doi:10.1007/s10641-017-0613-2

Waples, R. K., Larson, W. A., & Waples, R. S. (2016). Estimating contemporary effective population size in non-model species using linkage disequilibrium across thousands of loci. Heredity, 117(4), 233–240. doi:10.1038/hdy.2016.60

Waples, R. S., Antao, T., & Luikart, G. (2014). Effects of Overlapping Generations on Linkage Disequilibrium Estimates of Effective Population Size. Genetics, 197(2), 769–780. doi:10.1534/genetics.114.164822

Waples, R. S., & Do, C. (2010). Linkage disequilibrium estimates of contemporary Ne using highly variable genetic markers: A largely untapped resource for applied conservation and evolution. Evolutionary Applications, 3(3), 244–262. doi:10.1111/j.1752-4571.2009.00104.x

Waples, R. S., & Gaggiotti, O. (2006). What is a population? An empirical evaluation of some genetic methods for identifying the number of gene pools and their degree of connectivity. Molecular Ecology, 15(6), 1419–1439. doi:10.1111/j.1365-294X.2006.02890.x

Weir, B. S., & Cockerham, C. C. (1984). Estimating F-Statistics for the analysis of population structure. Evolution, 38(6), 1358–1370. doi:10.2307/2408641

Wellenreuther, M., & Bernatchez, L. (2018). Eco-evolutionary genomics of chromosomal inversions. Trends in Ecology and Evolution.

Whitlock, M. C. (2015). Modern Approaches to Local Adaptation. The American Naturalist, 186(S1), S1–S4. doi:10.1086/682933

Whitlock, M. C., & Lotterhos, K. E. (2015). Reliable Detection of Loci Responsible for Local Adaptation: Inference of a Null Model through Trimming the Distribution of F ST. The American Naturalist, 186(S1), S24–S36. doi:10.1086/682949

