## Supplementary material for "Chromosomal fusion and life history-associated genomic variation contribute to within-river local adaptation of Atlantic salmon"

Table S1: Summary of Miramichi River watershed Atlantic salmon sampling sites.

| Tributary | Trib. Code | Site | N | Date | Lat | Long |
| --- | --- | --- | --- | --- | --- | --- |
| Bartholomew River | BAR | BAR_Barth | 23 | 2016-10-07 | 46.6703 | -66.0399 |
|  |  | BAR_no119 | 2 | 2016-09-20 | 46.7196 | -65.8701 |
| Burnthill Brook | BHB | BHB_BH | 24 | 2016-10-06 | 46.6007 | -66.8385 |
|  |  | BHB_no120 | 2 | 2016-09-26 | 46.6814 | -66.9700 |
| Cains River | CAN | CAN_no110 | 8 | 2016-09-19 | 46.5327 | -65.8554 |
|  |  | CAN_no212 | 7 | 2016-09-12 | 46.3186 | -66.2893 |
|  |  | CAN_no74 | 10 | 2016-09-08 | 46.5828 | -65.7208 |
|  |  | CAN_no77 | 3 | 2016-09-12 | 46.4048 | -66.1094 |
|  |  | CAN_no78 | 14 | 2016-09-12 | 46.4349 | -66.0172 |
|  |  | CAN_Sabies | 15 | 2016-10-05 | 46.5858 | -65.7215 |
| Clearwater Brook | CWB | CWB_CW2 | 16 | 2016-10-06 | 46.6548 | -66.7688 |
|  |  | CWB_no121 | 6 | 2016-09-26 | 46.7572 | -66.8277 |
| Dungarvon River | DUN | DUN_no117 | 19 | 2016-10-03 | 46.7827 | -65.9729 |
|  |  | DUN_no186 | 12 | 2016-09-22 | 46.8209 | -66.6425 |
|  |  | DUN_no210 | 7 | 2016-09-22 | 46.7010 | -66.4839 |
|  |  | DUN_no55 | 6 | 2016-09-27 | 46.6590 | -66.3226 |
|  |  | DUN_no57 | 5 | 2016-09-27 | 46.6590 | -66.3226 |
| Lower Northwest | LNW | LNW_no215 | 4 | 2016-08-31 | 47.1350 | -65.8325 |
|  |  | LNW_no23 | 13 | 2016-09-02 | 46.9381 | -65.8217 |
|  |  | LNW_no26 | 1 | 2016-09-06 | 47.0550 | -65.8324 |
|  |  | LNW_no30 | 6 | 2016-08-31 | 47.2101 | -65.8178 |
|  |  | LNW_no40 | 10 | 2016-09-06 | 47.0420 | -65.8889 |
| Little Southwest | LSW | LSW_CatBr1 | 4 | 2016-10-11 | 46.8777 | -66.1082 |
|  |  | LSW_CatBr2 | 11 | 2016-10-13 | 46.8798 | -66.1025 |
|  |  | LSW_no107 | 2 | 2016-09-21 | 46.9650 | -66.6217 |
|  |  | LSW_no145 | 4 | 2016-09-21 | 46.9830 | -66.5168 |
|  |  | LSW_no147 | 21 | 2016-09-01 | 46.9821 | -66.3873 |
|  |  | LSW_no218 | 12 | 2016-09-21 | 46.9180 | -66.2895 |
|  |  | LSW_no43 | 1 | 2016-09-06 | 46.9369 | -65.9082 |
|  |  | LSW_Otter | 28 | 2016-10-13 | 46.8801 | -66.0355 |
| Main Southwest | MSW | MSW_Don | 24 | 2016-10-07 | 46.6044 | -65.8905 |
|  |  | MSW_no58 | 2 | 2016-09-30 | 46.4839 | -66.4819 |
|  |  | MSW_no61 | 1 | 2016-09-19 | 46.6088 | -65.8922 |
|  |  | MSW_no79_82 | 15 | 2016-09-08 | 46.5387 | -66.1868 |
|  |  | MSW_no84 | 4 | 2016-09-29 | 46.4607 | -66.4085 |
|  |  | MSW_Porter | 10 | 2016-10-05 | 46.4820 | -66.4733 |
| Northwest Millstream | NWM | NWM_BB | 4 | 2016-10-12 | 47.1086 | -65.6177 |
|  |  | NWM_OxB | 17 | 2016-10-12 | 47.1184 | -65.6478 |
|  |  | NWM_OxBrte430 | 26 | 2016-10-12 | 47.0464 | -65.6395 |
| Rocky Brook | RBR | RBR_no92 | 21 | 2016-09-30 | 46.6892 | -66.6358 |
|  |  | RBR_rbr | 25 | 2016-10-06 | 46.6845 | -66.6328 |
| Renous River | REN | REN_no214 | 4 | 2016-10-03 | 46.7908 | -66.4764 |
|  |  | REN_no48_116 | 13 | 2016-09-27 | 46.8055 | -65.8715 |
|  |  | REN_no54 | 8 | 2016-09-22 | 46.7968 | -66.2017 |
|  |  | REN_rte108 | 23 | 2016-10-12 | 46.7933 | -66.1968 |
| Sevogle River | SEV | SEV_no103 | 12 | 2016-09-06 | 47.0519 | -65.9987 |
|  |  | SEV_no104 | 6 | 2016-08-29 | 47.1352 | -65.9821 |
|  |  | SEV_no153 | 10 | 2016-09-01 | 47.0903 | -66.3177 |
|  |  | SEV_no38 | 10 | 2016-08-31 | 47.1327 | -66.1182 |
|  |  | SEV_no39 | 10 | 2016-09-01 | 47.0688 | -66.2589 |
| Taxis River | TAX | TAX_no86 | 7 | 2016-09-29 | 46.4311 | -66.5793 |
|  |  | TAX_tax | 14 | 2016-10-05 | 46.4255 | -66.6055 |
|  |  | TAX_trib | 23 | 2016-10-05 | 46.4329 | -66.5794 |
| Upper Northwest | UNW | UNW_no115 | 22 | 2016-08-29 | 47.1895 | -65.8923 |
|  |  | UNW_no20 | 6 | 2016-08-29 | 47.1918 | -65.9076 |
|  |  | UNW_no31 | 12 | 2016-08-29 | 47.1669 | -65.9556 |
|  |  | UNW_no33 | 7 | 2016-08-31 | 47.1909 | -66.1210 |
|  |  | UNW_no34 | 14 | 2016-08-30 | 47.2531 | -66.2362 |
|  |  | UNW_no35 | 16 | 2016-08-30 | 47.2721 | -66.3213 |
| Upper Southwest | USW | USW_Mclean | 21 | 2016-10-06 | 46.5537 | -66.8563 |
|  |  | USW_no129 | 12 | 2016-09-26 | 46.5492 | -67.0435 |
|  |  | USW_no206 | 11 | 2016-09-28 | 46.5619 | -67.2901 |
|  |  | USW_no60 | 15 | 2016-09-30 | 46.5968 | -66.6913 |
|  |  | USW_no69 | 7 | 2016-09-28 | 46.5110 | -67.1130 |

Trib. Code: Tributary population code used throughout the manuscript and figures, N: sample size, Lat: Latitude (decimal degrees), Long: Longitude (decimal degrees).

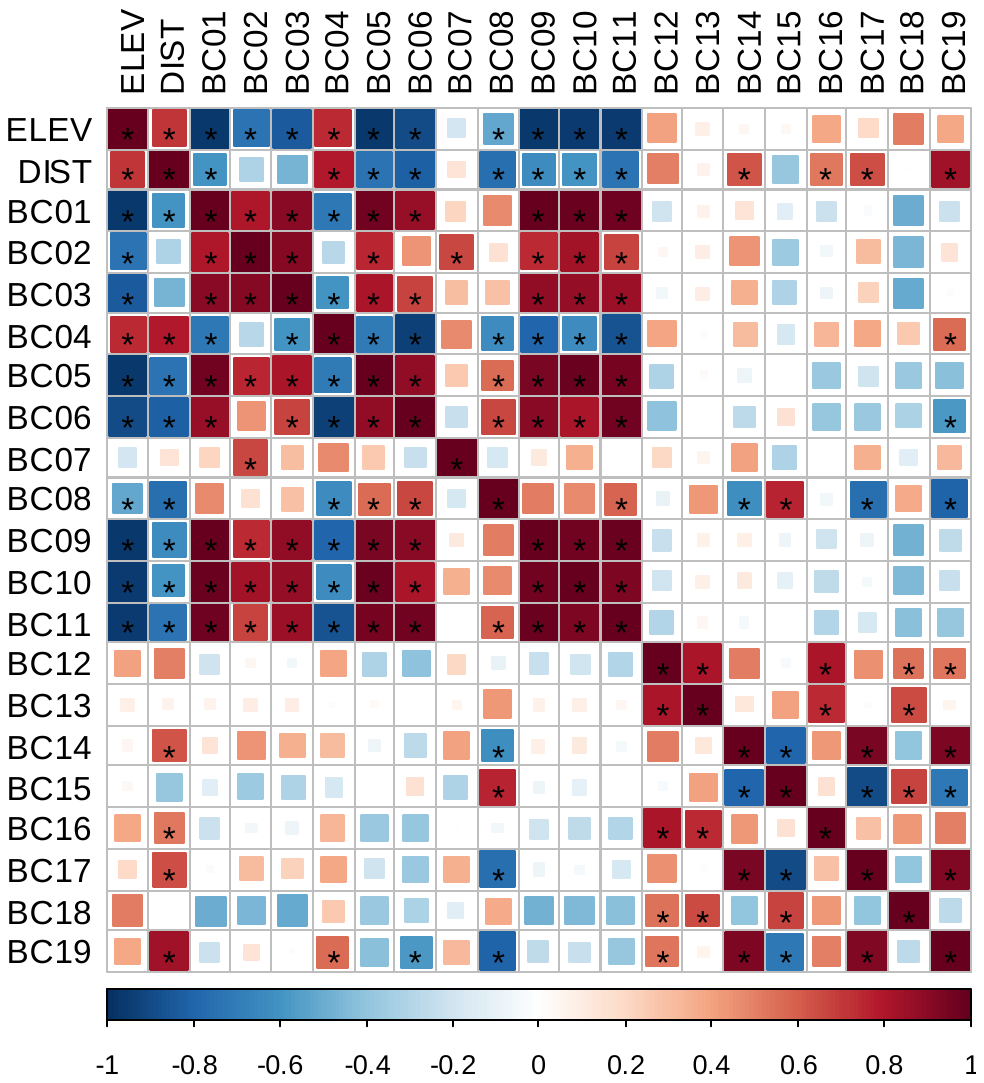

Figure S1: Correlations among environmental variables extracted from the WorldClim 2.0 Bioclimatic database (http://www.worldclim.org/bioclim). Colors indicate direction (red: positive, blue: negative) and magnitude of Pearson’s correlation coefficient (r) and asterisks indicate statistical significance of the relationship (p < 0.05). ELEV = elevation above sea level, DIST = distance upstream from the river mouth, BC01 = annual mean temperature, BC2 = mean diurnal range (mean of monthly (max. temp – min. temp)), BC03 = isothermality (BC02/BC07 * 100), BC04 = temperature seasonality (standard deviation *100), BC05 = max. temperature of warmest month, BC06 = min. temperature of coldest month, BC07 = temperature annual range (BC05-BC06), BC08 = mean temperature of wettest quarter, BC09 = mean temperature of driest quarter, BC10 = mean temperature of warmest quarter, BC11 = mean temperature of coldest quarter, BC12 = annual precipitation, BC13 = precipitation of wettest month, BC14 = precipitation of driest month, BC15 = precipitation seasonality (coefficient of variation), BC16 = precipitation of wettest quarter, BC17 = precipitation of driest quarter, BC18 = precipitation of warmest quarter, BC19 = precipitation of coldest quarter.

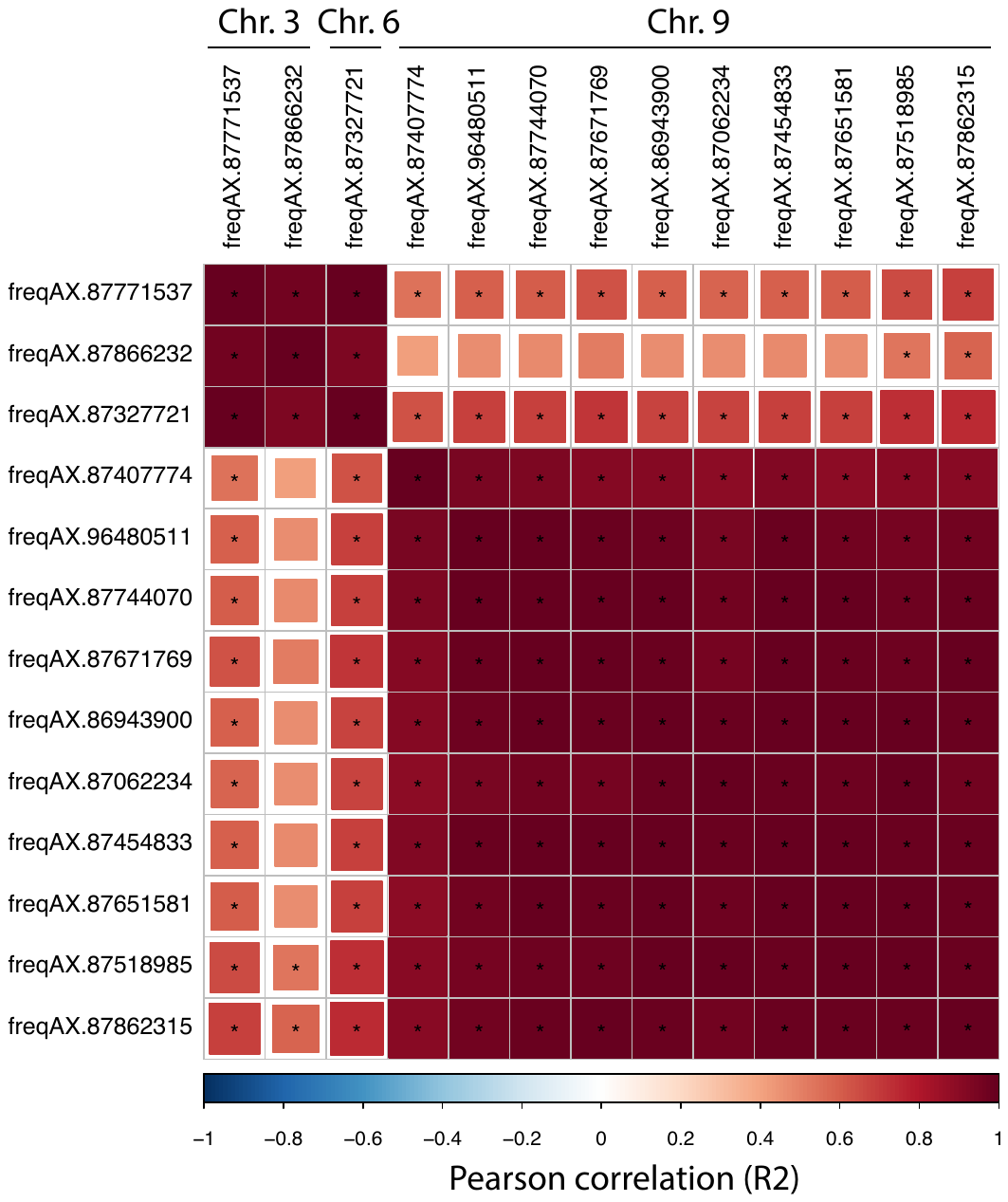

Figure S2: Correlations among minor allele frequencies for 13 outlier SNPs detected by all single locus outlier tests. Colors indicate direction (red: positive, blue: negative) and magnitude of Pearson’s correlation coefficient (r) and asterisks indicate statistical significance of the relationship (p < 0.05).

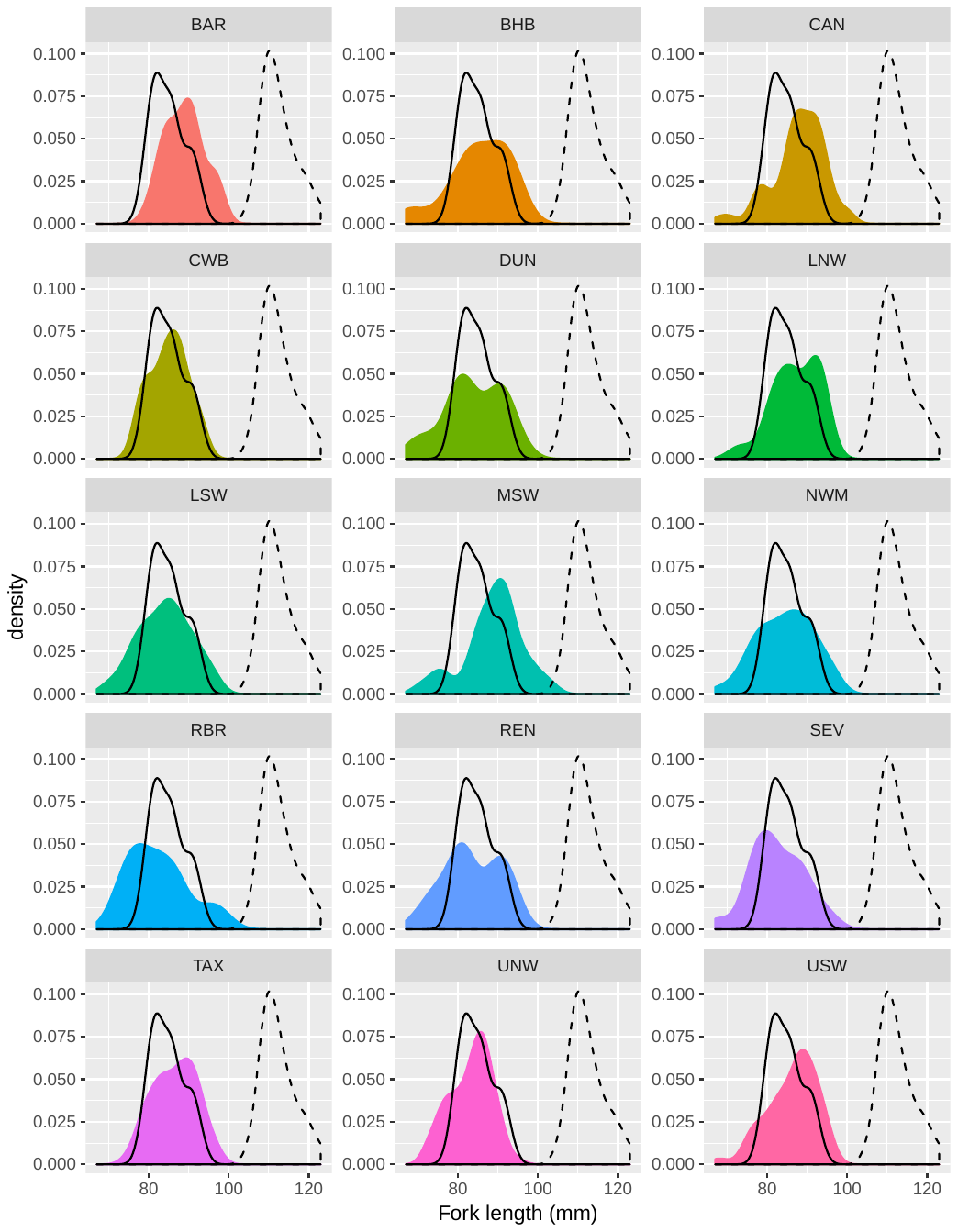

Figure S3: Frequency distributions of 1+ age Atlantic salmon fork length from 15 major tributaries of the Miramichi River. Solid black and dashed lines represent distribution of annual mean lengths of 1+ and 2+ age Atlantic salmon respectively (Means for years 1970 – 2000: adapted from Swansburg *et al.* 2002). Note: the distributions of annual means presented underestimate the variability in size distributions for individuals in a given year, but do provide a useful approximation for estimating the general size distinction between age 1+ and 2+ juvenile salmon in this river.

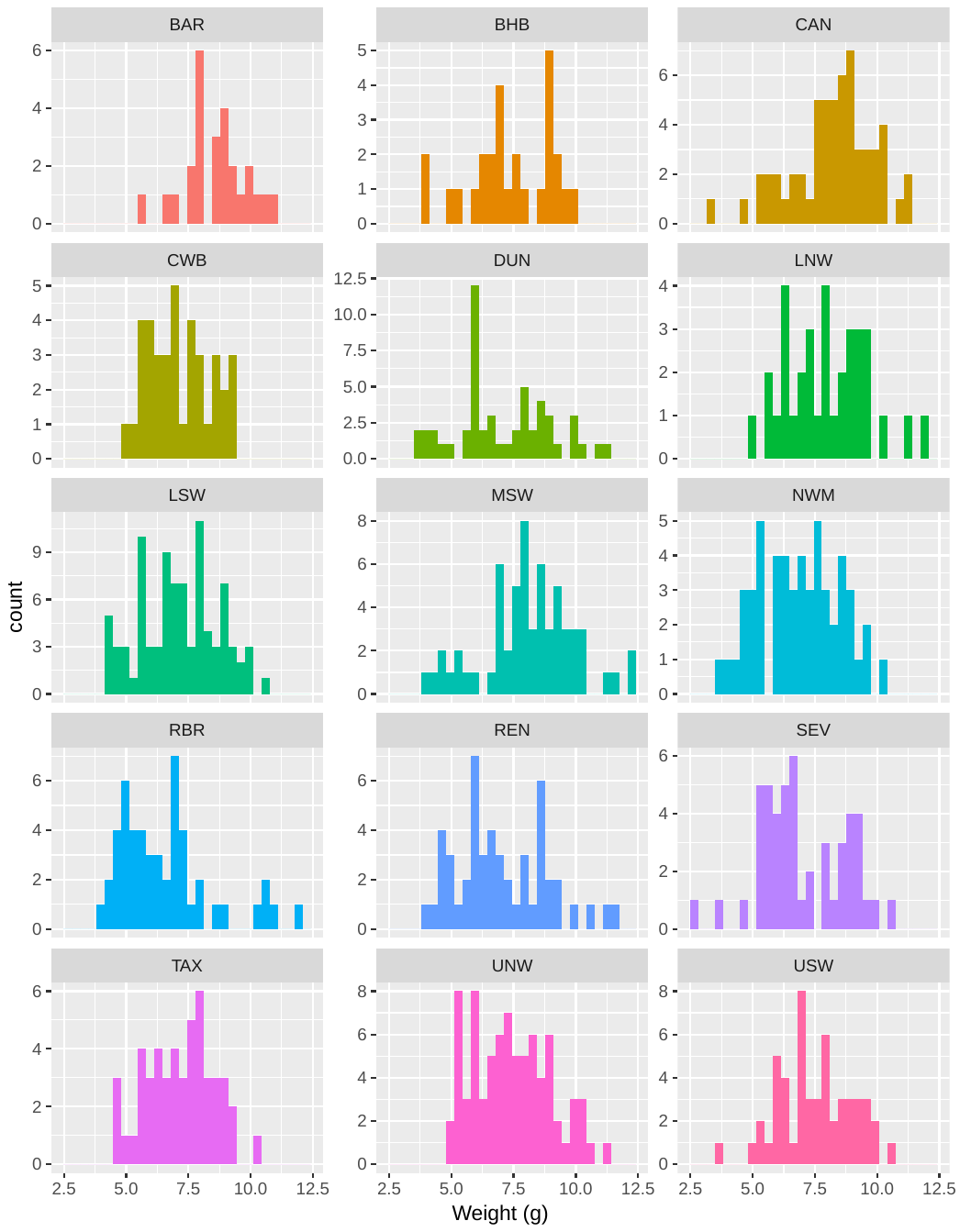

Figure S4: Frequency distributions for weight of 1+ age Atlantic salmon from 15 major tributaries of the Miramichi River.

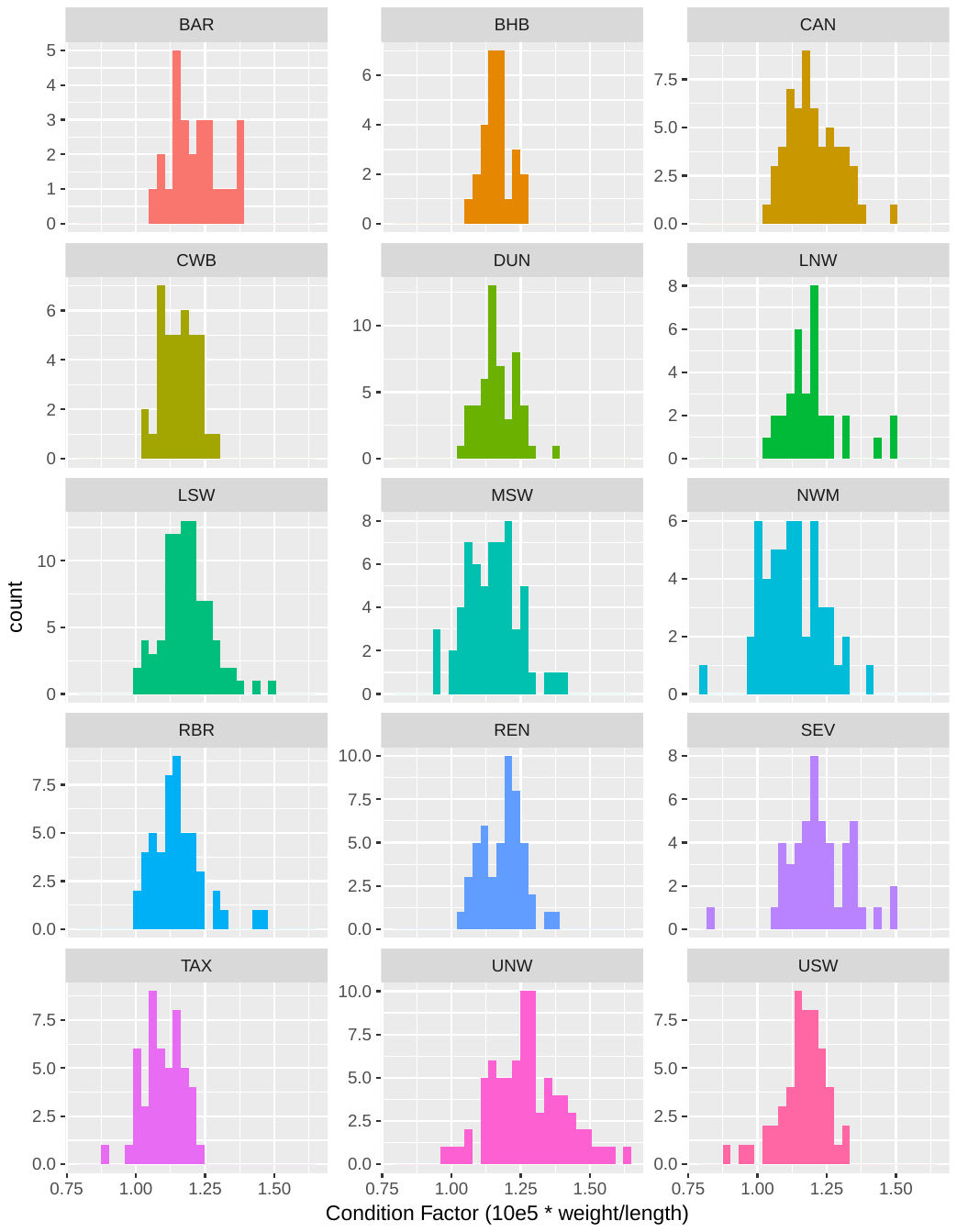

Figure S5: Frequency distributions for condition factor of 1+ age Atlantic salmon from 15 major tributaries of the Miramichi River.

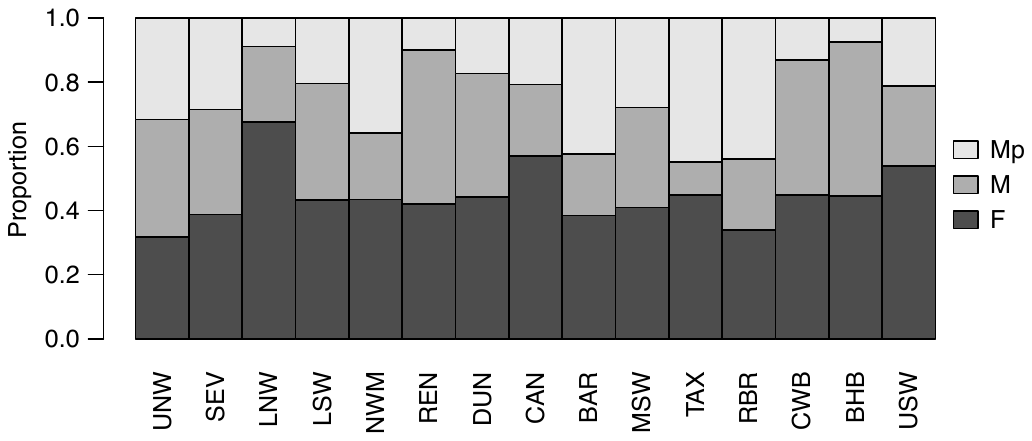

Figure S6: Sex proportions (F = females, M = males, Mp = precociously mature males) of 1+ juvenile Atlantic salmon from 15 tributary populations within the Miramichi River. See Table 1 for tributary code descriptions.

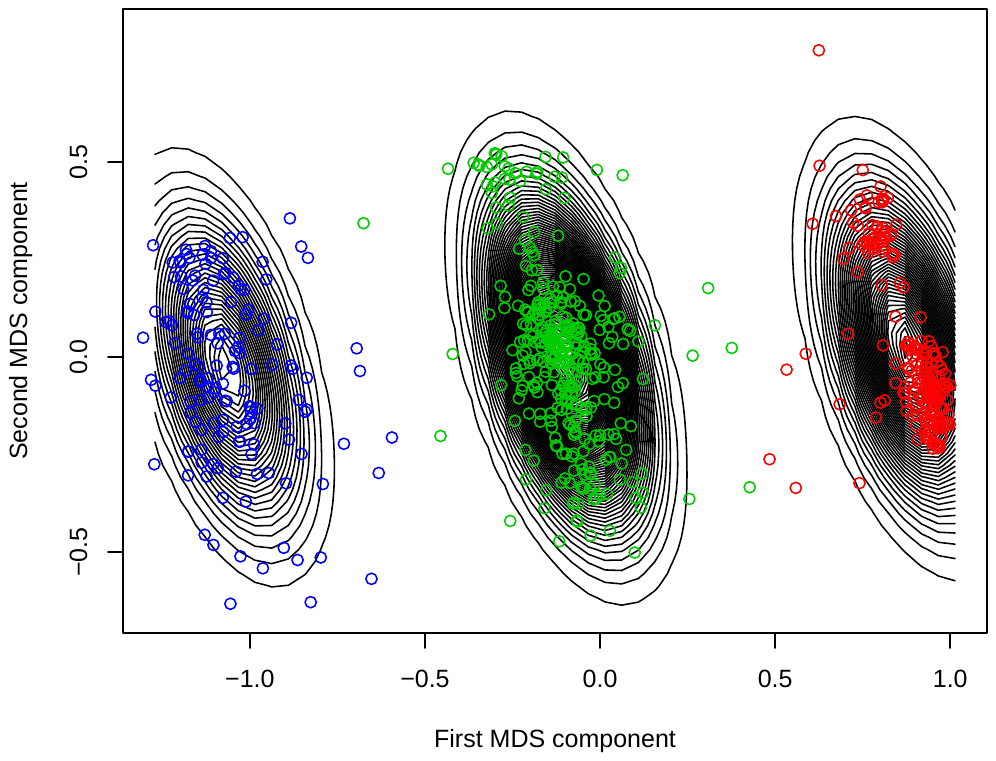

Figure S7: InvClust PCA analysis of the highly linked regions of chromosome 8 and 29 for assigning fusion karyotype to individuals.

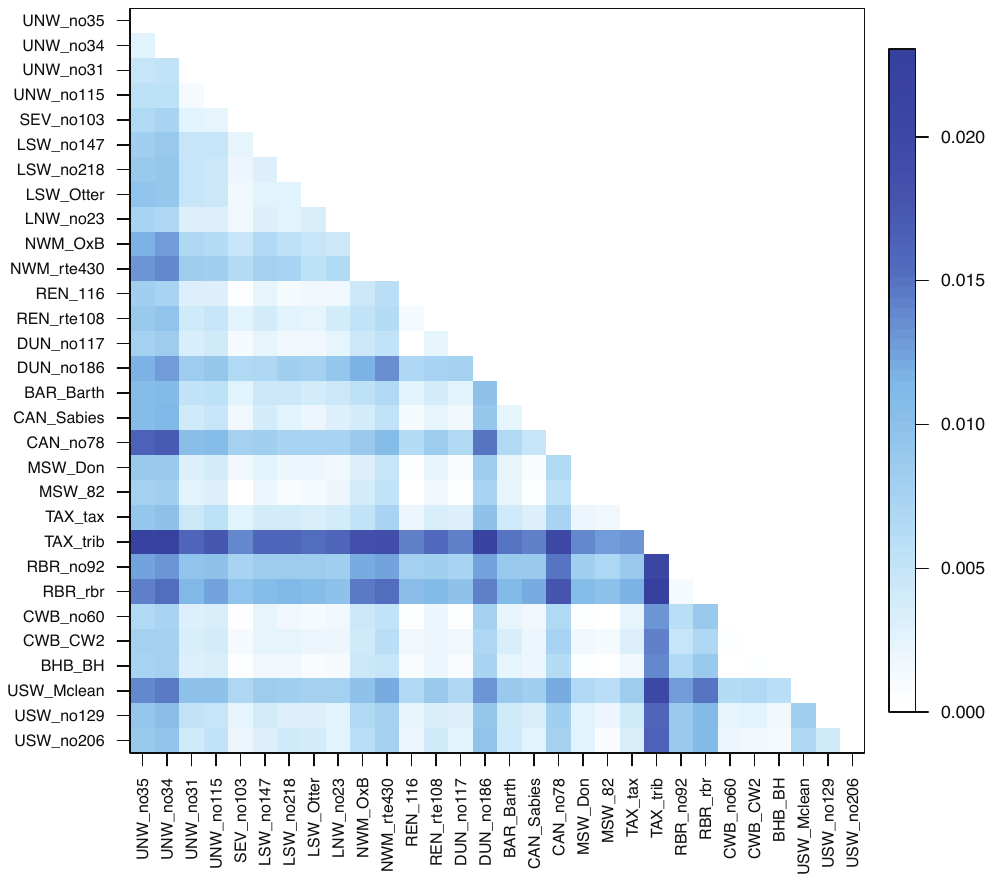

Figure S8: Pairwise F_ST_ among sites possessing greater than 10 individuals.

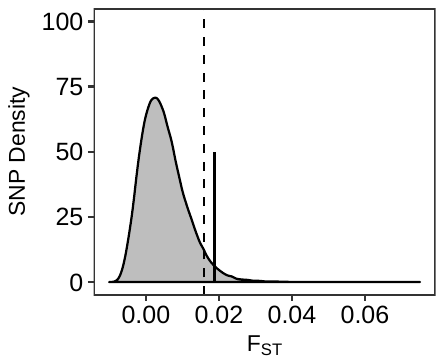

Figure S9: Distribution of global F_ST_ among tributary populations of Atlantic salmon in the Miramichi River based on 52466 SNPs. The divergence among tributaries based on chromosome 8-29 fusion karyotype is indicated by the solid line and the 95^th^ percentile of the SNP distribution is indicated by the dashed line.

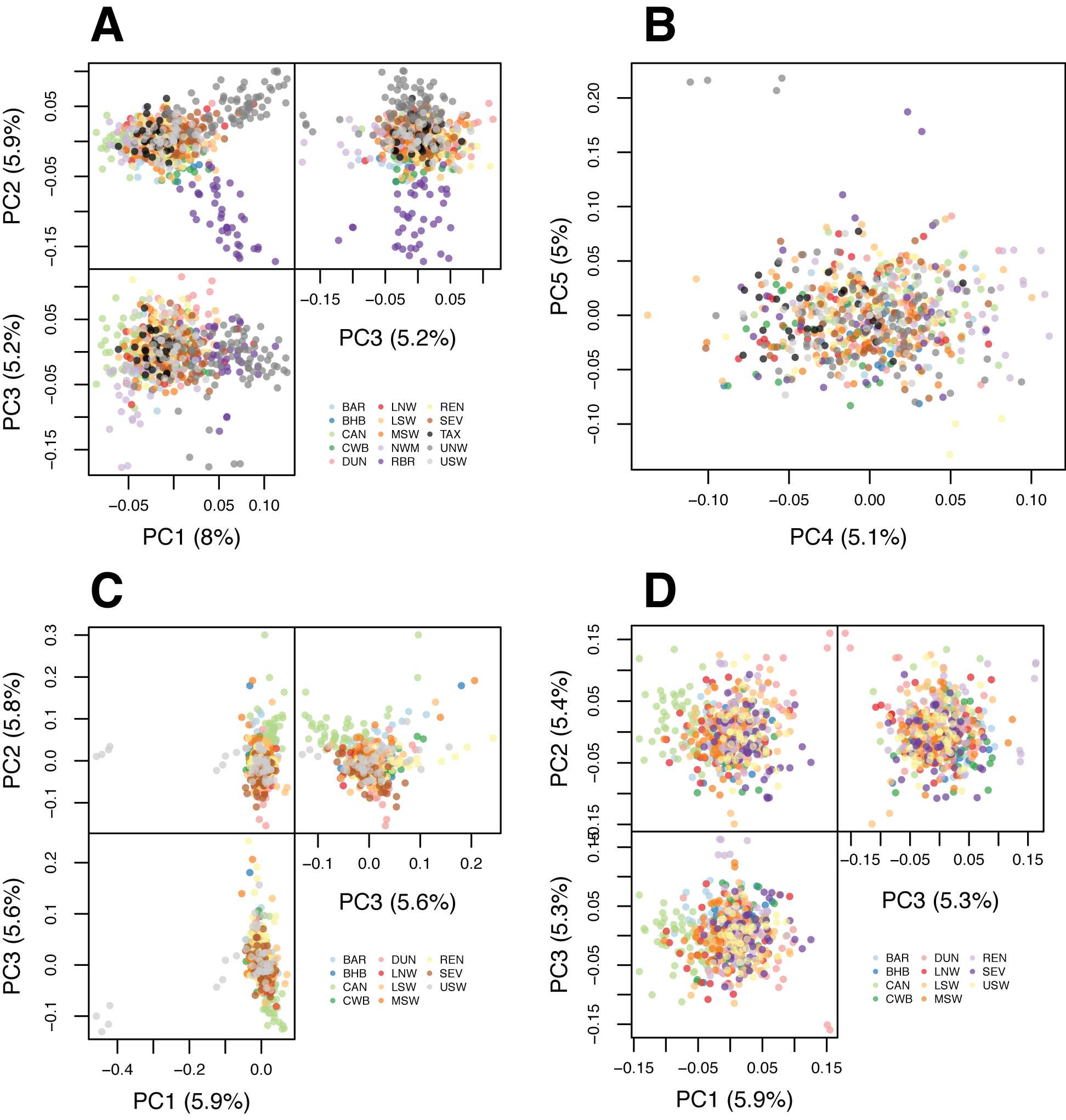

Figure S10: Supplemental principal components analysis of putatively neutral SNP data. A) PCA conducted using EIGENSOFT to exclude outlier samples, B) Axes 4 and 5 of EIGENSOFT PCA, C) PCA following removal of divergent tributaries (NWM, UNW, TAX, RBR) implemented in PLINK and D) PCA following removal of divergent tributaries (NWM, UNW, TAX, RBR) implemented in EIGENSOFT.

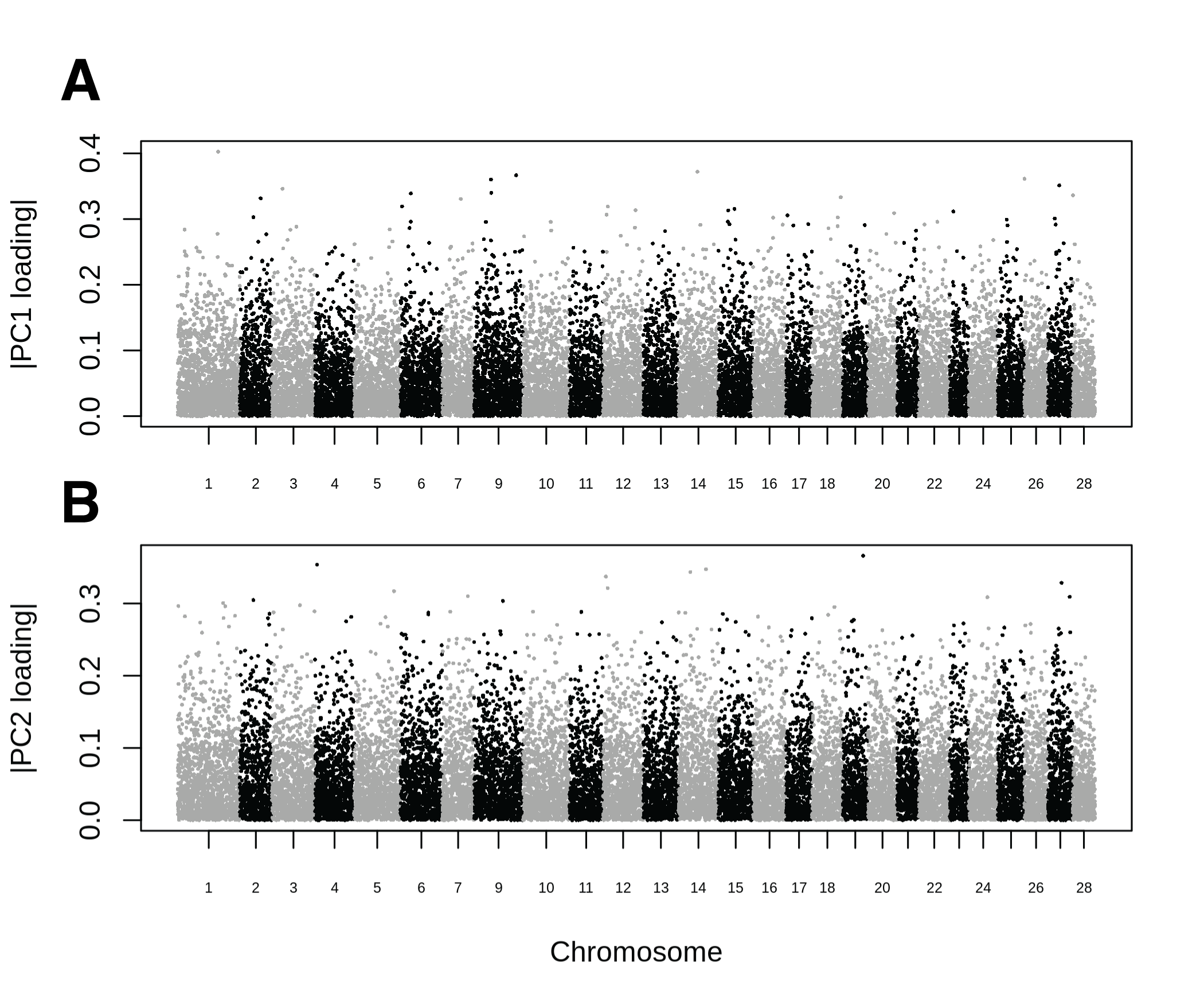

Figure S11: Absolute value of SNP PC loadings for PC axes 1 (A) and 2 (B) from PCA using neutral SNPs and all 15 tributary populations of Atlantic salmon from the Miramichi River.

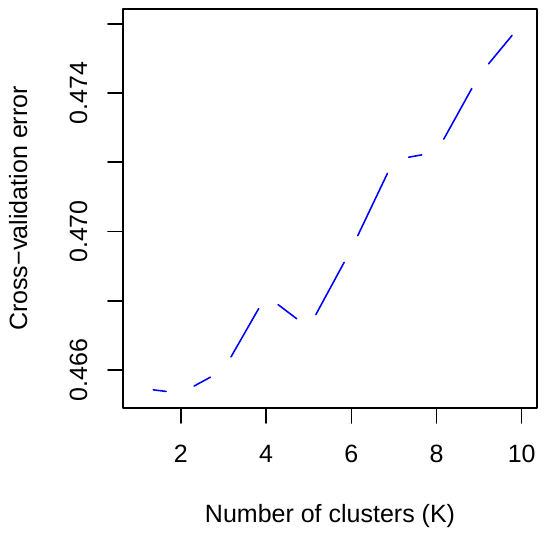

Figure S12: Admixture cross-validation error (5-fold) for K = 1 to K = 10.

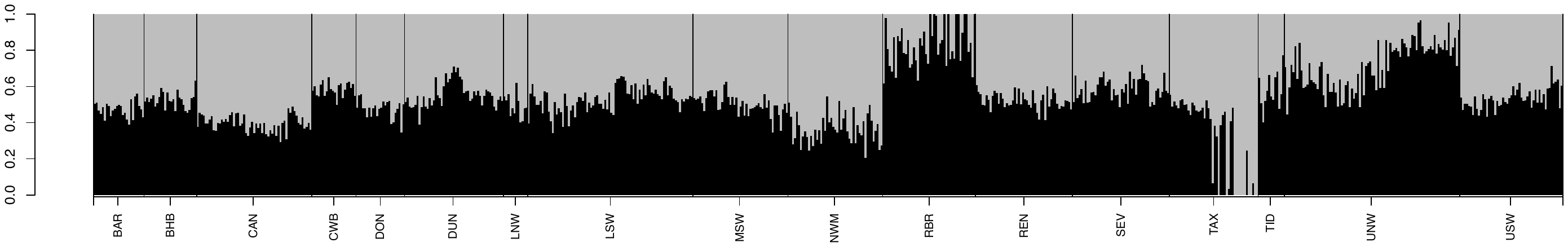

Figure S13: Admixture coefficients for 728 Atlantic salmon at K = 2 based on 22105 SNPs. Given the weak support for K=2 based on cross-validation analysis this figure should not be interpreted literally in the sense that our samples represent genetic admixture of two ancestral populations. We provide this figure to emphasize the lack of structure based on naïve clustering routines. The appearance pure “grey” individuals in TAX likely reflects a demographic bottleneck experienced by this population, that is also reflected in the reduced effective number of breeders, rather than representing a true ancestral population as is assumed by the software.

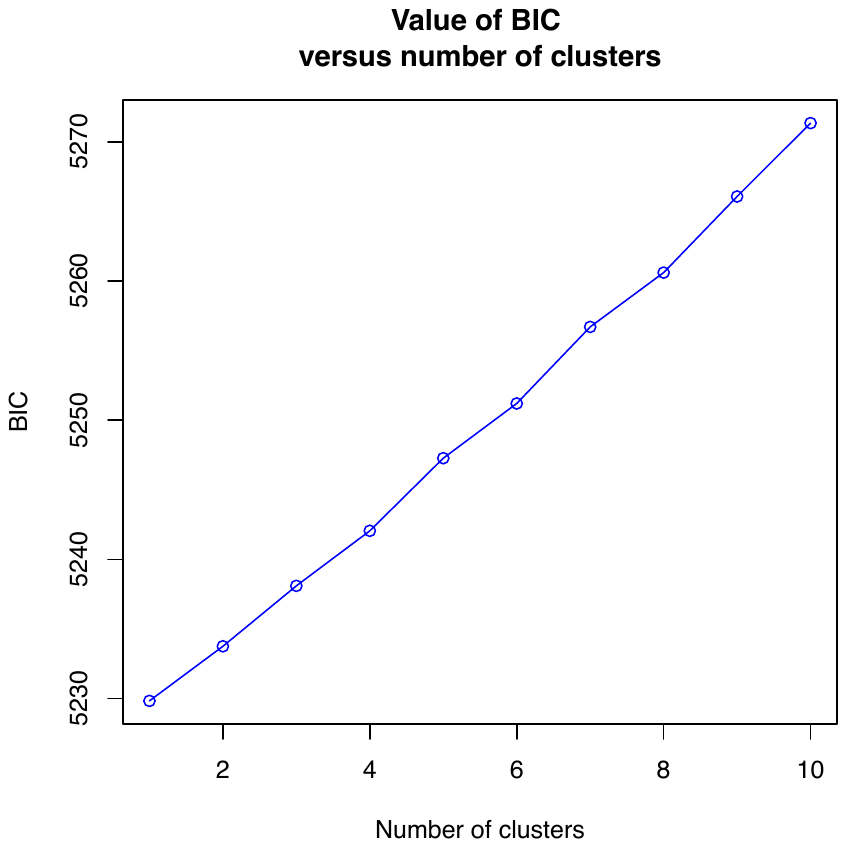

Figure S14: Bayesian Information Criterion for k-means clustering of principal component summarized allelic variation number of genetic clusters from K = 1 to K = 10.

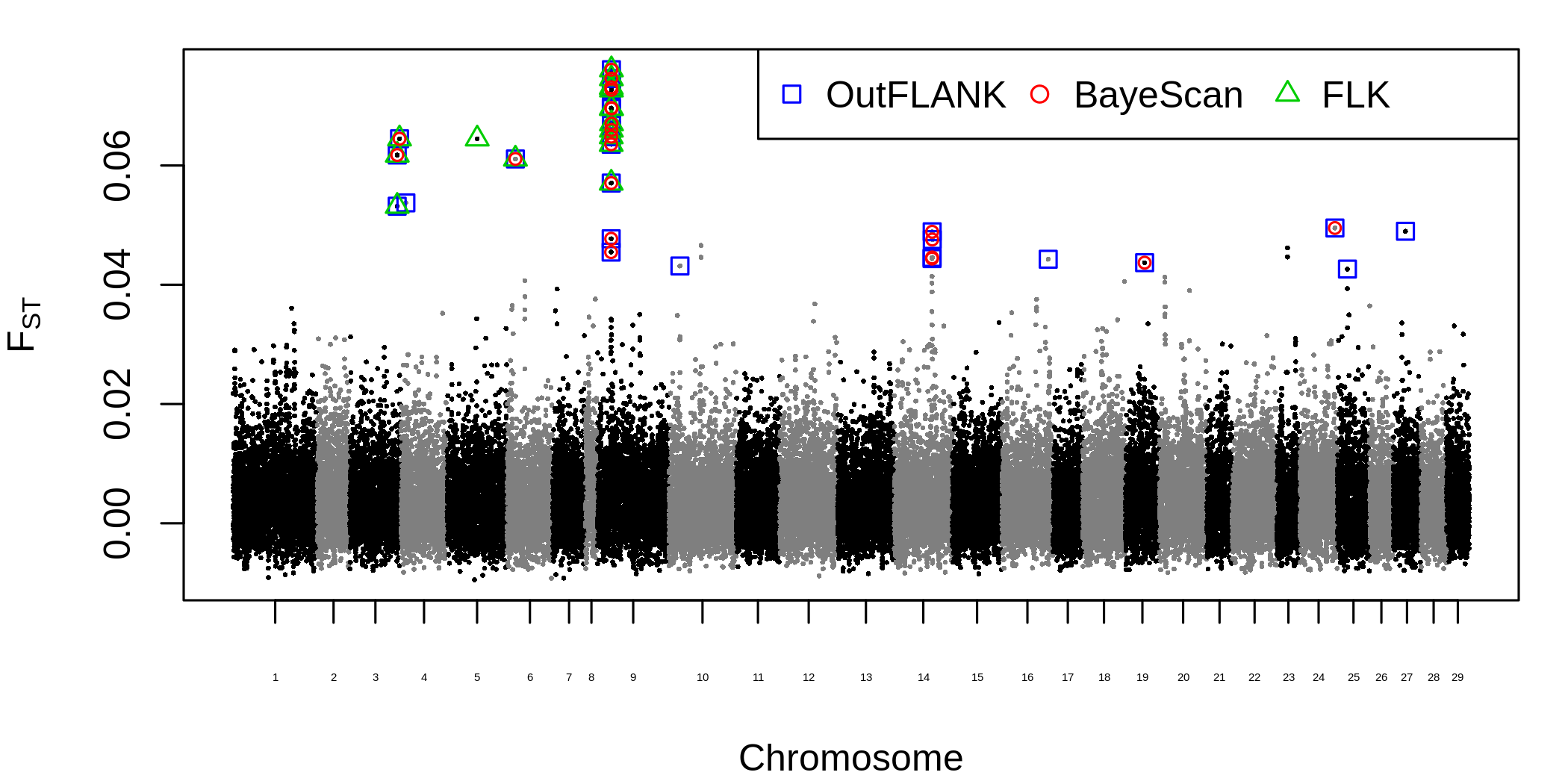

Figure S15: Genome-wide distribution of F_ST_ outliers using three different outlier detection methods.

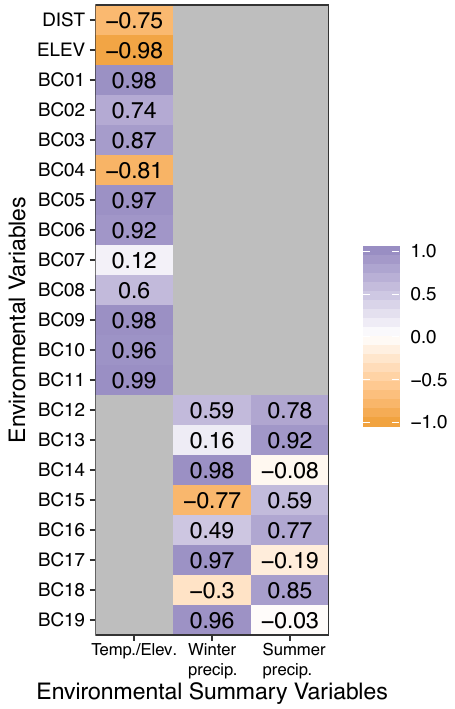

Figure S16: PCA loadings (Pearson correlations) of the WorldClim 2.0 (http://worldclim.org/bioclim) bioclimatic variables (BC01-19), elevation (ELEV), and distance from the ocean (DIST) on the three retained environmental summary variables.

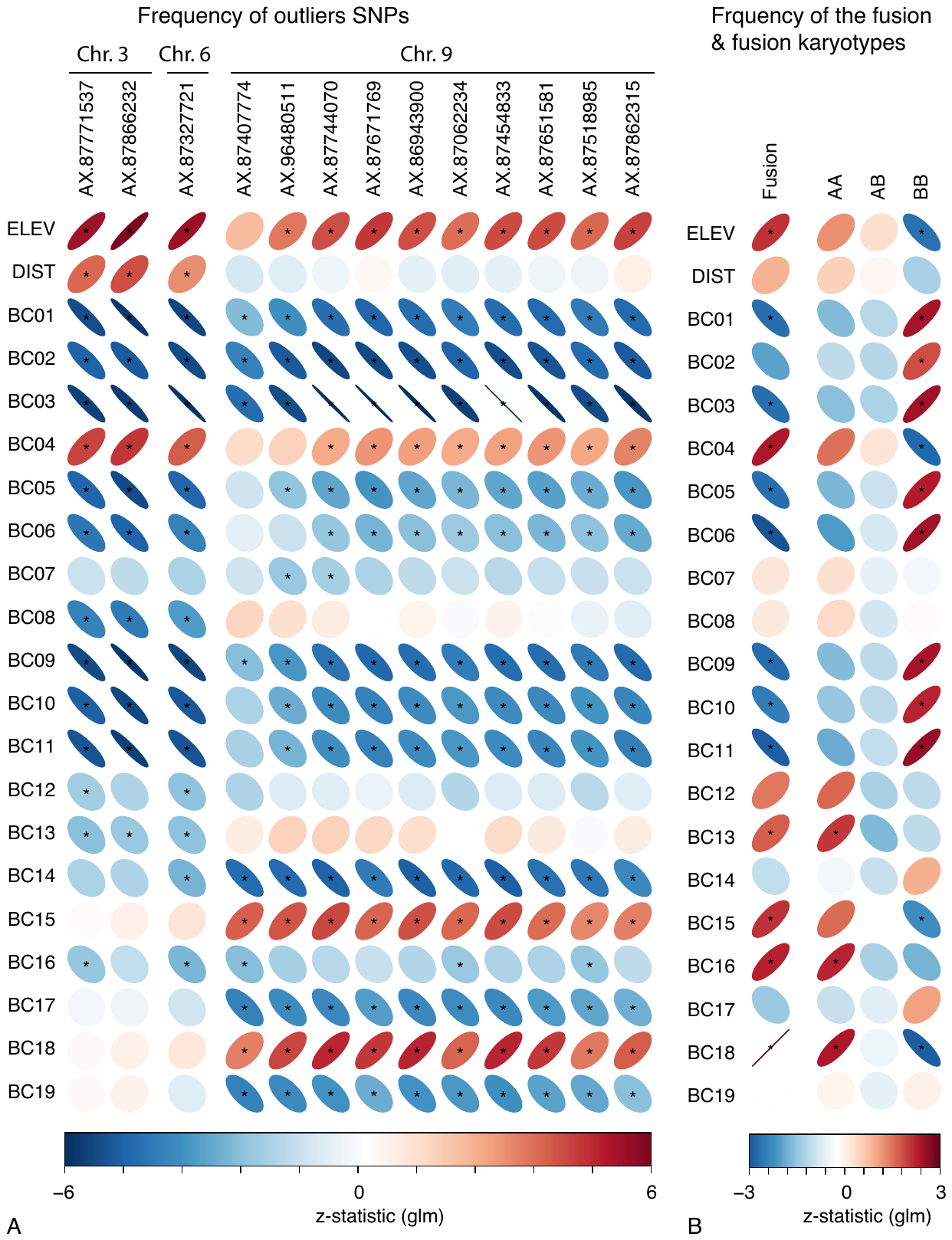

Figure S17: Correlations of minor allele frequencies for 13 outlier SNPs (A) and fusion and karyotype frequency (B) with individual temperature and precipitation variables from the WorldClim 2.0 bioclimatic database. Strength and direction of the statistical association (GLM z) are indicated by the shape of the ellipse and its colour (red: positive, blue: negative). Stars denote significance at FDR = 0.05 level (Benjamini and Hochberg 1995).

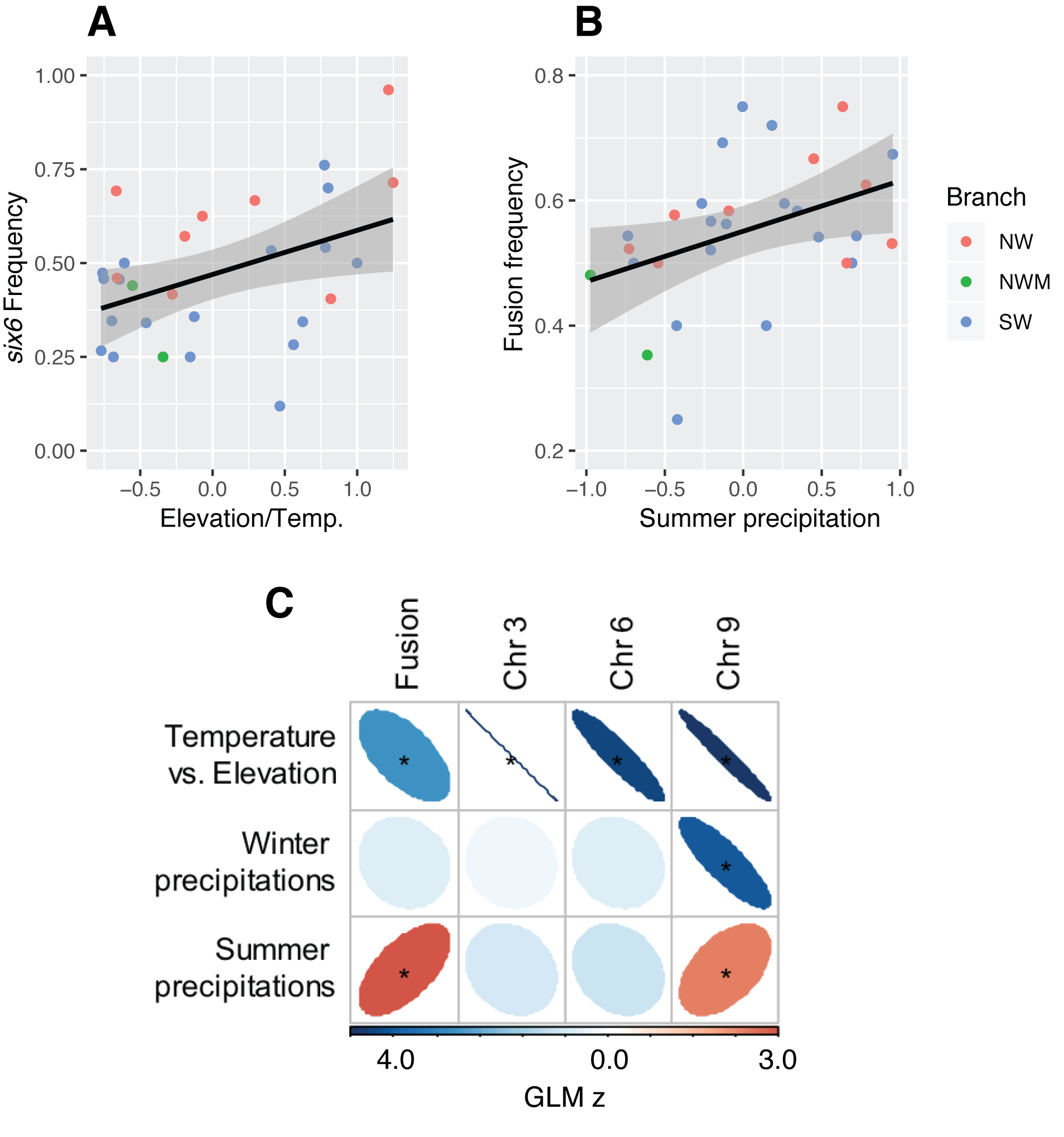

Figure S18: Genomic variation associated with environmental attributes at the site level for sites with >=12 individuals. (A) six6 region of chromosome 9 with elevation/temp. gradient (GLM, z = -4.0, p < 0.001), (B) fusion frequency correlation with summer precipitation (GLM, z = 3.0, p = 0.06), and (C) all associations.
